# Improving gene regulatory network inference and assessment: The importance of using network structure

**DOI:** 10.1101/2023.01.12.523840

**Authors:** Juan M. Escorcia-Rodríguez, Estefani Gaytan-Nuñez, Ericka M. Hernandez-Benitez, Andrea Zorro-Aranda, Marco A. Tello-Palencia, Julio A. Freyre-González

## Abstract

Gene regulatory networks are graph models representing cellular transcription events. Networks are far from complete due to time and resource consumption for experimental validation and curation of the interactions. Previous assessments have shown the modest performance of the available network inference methods based on gene expression data. Here, we study several caveats on the inference of regulatory networks and methods assessment through the quality of the input data and gold standard, and the assessment approach with a focus on the global structure of the network. We used synthetic and biological data for the predictions and experimentally-validated biological networks as the gold standard (ground truth). Standard performance metrics and graph structural properties suggest that methods inferring co-expression networks should no longer be assessed equally with those inferring regulatory interactions. While methods inferring regulatory interactions perform better in global regulatory network inference than co-expression-based methods, the latter is better suited to infer function-specific regulons and co-regulation networks. When merging expression data, the size increase should outweigh the noise inclusion and graph structure should be considered when integrating the inferences. We conclude with guidelines to take advantage of inference methods and their assessment based on the applications and available expression datasets.

## Introduction

A gene regulatory network (GRN) is responsible for sensing environmental cues and responding accordingly. It represents directed regulatory interactions between genes coding transcription factors (TFs) and their target genes (TGs). Successful developments in synthetic biology require that the designed circuit properly integrates into the global and local regulatory circuits (Freyre-Gonzalez et al., 2022). This is a current challenge as there is not a single complete experimentally-validated GRN (Escorcia-Rodriguez et al., 2020), only a handful (< 4) of bacterial organisms has a known GRN having completeness > 70%, and its experimental reconstruction is a time- and resource-consuming task. Consequently, computational network inference is frequently used. Whereas previous works have evaluated network inference tools using synthetic and experimental data for several organisms (Marbach et al., 2010; Marbach et al., 2012; Chen and Mar, 2018), they did not assess several essential criteria for the inference of GRNs such as data noise variation, and the global structure of the predictions and the gold standard (GS). Riet De Smet and Kathleen Marchal reviewed the advantages and limitations of several inference methods through the biological interpretation of the network structure but did not use the structure itself to assess the predictions (De Smet and Marchal, 2010).

Employing artificial data with varying amounts of noise, Deniz Seçilmiş et al. recently evaluated various tools and discovered that using the perturbation design matrix outperformed methods without it. (Secilmis et al., 2022). Synthetic data are the first alternative for benchmarking inference methods (Van den Bulcke et al., 2006). However, the generation of synthetic data relies on simulation parameters (e.g., dimension and noise of the dataset), which may not reflect the variability in biological data. Regarding the transcriptomic technique, most of the tools developed for GRN inference from microarray data have been indiscriminately coupled with RNA-seq (Iancu et al., 2012; Salleh et al., 2018; Zhang et al., 2019) despite tools for bulk RNA-seq data have been already developed (Proost et al., 2017; Imbert et al., 2018).

The authors of the DREAM5 network inference challenge evaluated a plethora of genome-scale transcriptional regulatory network predictions from gene expression data. Their results provided insights into the difficulty of GRN inference using correlation and mutual information between gene pairs and found that contrary to synthetic data, the dependencies between genes interacting in the cell barely exceeded the dependencies between non-interacting gene pairs in biological data. Interestingly, with synthetic and *Escherichia coli* data, the correlations between genes regulated by identical sets of TFs exceeded those between genes in the actual regulatory network (Supplementary Note 5 in (Marbach et al., 2012)), but most of those interactions between co-regulated genes would be false positives (e.g., structural genes shaping a transcription unit). Recently, Simon Larsen et al. performed an in-deep analysis on this matter, their results show that the correlation of pairs of random genes is indistinguishable from those involved in known regulatory interactions in *E. coli* (Larsen et al., 2019). Doglas Parise et al. confirmed the results on *Corynebacterium glutamicum* (Parise et al., 2021).

According to the DREAM5 team, integrating predictions from different inference techniques through the Borda count method (“community network”) is the best strategy because method performance is not consistent across species. (Marbach et al., 2012). Since then, the community approach has been broadly applied (Akesson et al., 2021; Zorro-Aranda et al., 2022). ComHub is a pipeline for integrating predictions from various methods to rank regulators according to their average out-degree using gene expression. (Akesson et al., 2021). Recently we inferred a GRN for *Streptomyces coelicolor* and identified the global regulators applying the NDA (natural decomposition approach) (Freyre-Gonzalez et al., 2008; Freyre-Gonzalez et al., 2012) on the across-methods community network preserving only TF-TG interactions (Zorro-Aranda et al., 2022). However, some methods are better suited to particular global topological structures (Stolovitzky et al., 2009). Thus, the hubs may differ across methods and have different biological interpretations in each global network due to the inherently different structure.

The inferences are commonly assessed using standard performance metrics such as the area under the recall vs precision (AUPR) and true negative rate vs recall curves. These metrics rely heavily on the ranking of the interactions (Marbach et al., 2010). Based on the ranking scheme and the cutoff value, the global network will also have a different structure. For example, using the Pearson correlation coefficient with no post-processing step as the ranking score, co-regulated genes from the same transcription unit (TU) will be at the top of the prediction and the global network will be shaped by interactions between co-expressed genes. This would be a good co-regulation network, but it will be highly penalized if it is assessed against a GRN. The edges represent different biological associations (De Smet and Marchal, 2010); therefore, the networks have a different global structure and are better suited for different purposes (Michoel et al., 2009). However, the assessment and integration of inference methods designed for co-expression are still being directly used and compared with those inferring regulation (Marbach et al., 2012; Bellot et al., 2015; Pratapa et al., 2020; Secilmis et al., 2022).

We previously explored structural properties and systems-level components to analyze curated and inferred GRNs for *Streptomyces coelicolor* (Zorro-Aranda et al., 2022). Here, we focused on the factors influencing the inference of GRNs and their assessment. Mainly, the structural characteristics of the GS and the inferred networks, the quality of the input data and the GS, and the assessment strategy. Besides synthetic data with varying noise and completeness levels, we use biological data for *Escherichia coli, Bacillus subtilis*, and *Pseudomonas aeruginosa* along with their experimentally-validated GRNs (Escorcia-Rodriguez et al., 2020) as the GS. Because the networks used as GS are not complete, unknown actual interactions identified in the prediction will be misclassified as a false positive. To check whether our results will hold when the GS networks are complete, we used historical snapshots with different completeness levels and evidence (Escorcia-Rodriguez et al., 2020). Figure 1 summarizes the complete workflow.

**Figure 1.**
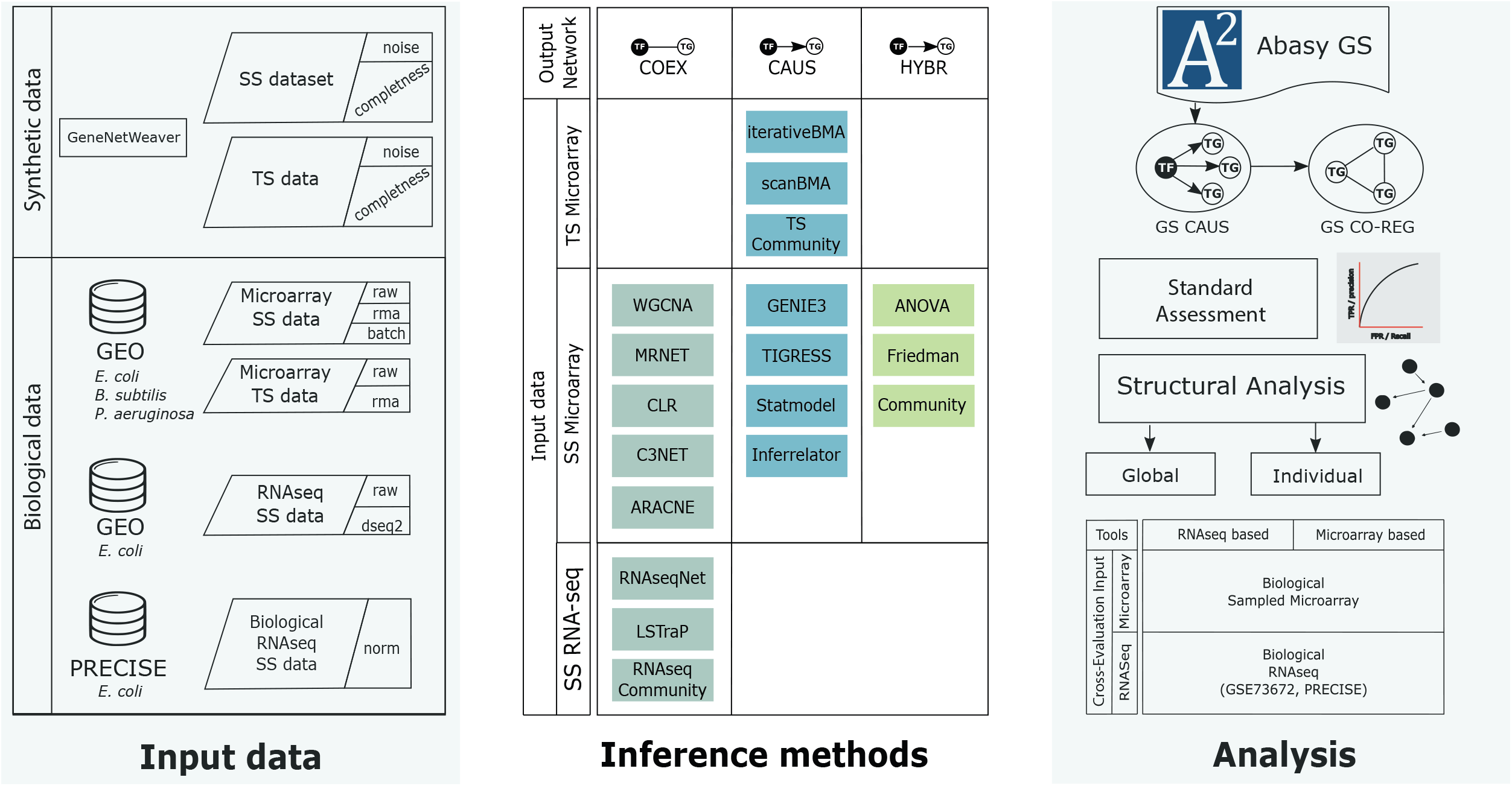
Workflow of this work. We generated synthetic data using GeneNetWeaver for *E. coli* and collected several biological microarray datasets from GEO for *E. coli, B. subtilis*, and *P. aeruginosa*, as well as RNA-seq data from GEO and PRECISE for *E. coli* (left column). The synthetic and biological datasets were used as input for the inference methods (middle row). The inference methods were classified according to their final network type. COEX tools generate undirected networks. CAUS tools generate directed networks using a list of regulators to compute the predictions as part of their algorithm. HYBR includes Friedman and ANOVA implementations (Zorro-Aranda et al., 2022) that generate co-expression networks that are trimmed to only include regulations mediated by a known transcription factor. The Community networks are classified according to the type of tools they include. We used biological networks as the gold standard to perform the assessment and analyses (right column). From the directed gold standard (“CAUS” GS) we generated a co-regulation gold standard (GS CO-REG). We performed the standard statistical and a structure-based assessment. **SS:** steady-state data, TS: time-series data, GS: gold standard, TF: transcription factor, TG: target gene. See Supplementary Figure 1 for further details.

## Results and discussion

We reviewed the literature to construct a collection of network inference tools. After the application of filter criteria (see Materials and methods), 15 tools were selected to be assessed along with “Community” reconstructions integrating interactions from several tools. Then, we arranged the inference tools according to the output network type into three groups (Table 1 and Figure 1): 1) The COEX tools infer interactions between genes with correlated expression profiles. 2) The CAUS tools use a TFs list to infer regulatory interactions between the TFs and their TGs (i.e., GRNs) (Hecker et al., 2009). 3) The HYBR (hybrid) group contemplates ANOVA (Kuffner et al., 2012) and Friedman (Zorro-Aranda et al., 2022) which are based on analysis of variance and therefore do not infer causality. However, we used a list of TFs to keep only TF-TG interactions. The classification of Community relies on the type of interactions that it includes. It is considered HYBR when it integrates interactions from different network types, but it will be considered CAUS if it only integrates interactions from CAUS tools. Similarly, Community will be considered COEX if it only contains interactions from COEX tools. See Table 1 and the ΣBackground section in the supplementary material for a detailed description of the tools.

**Table 1.**
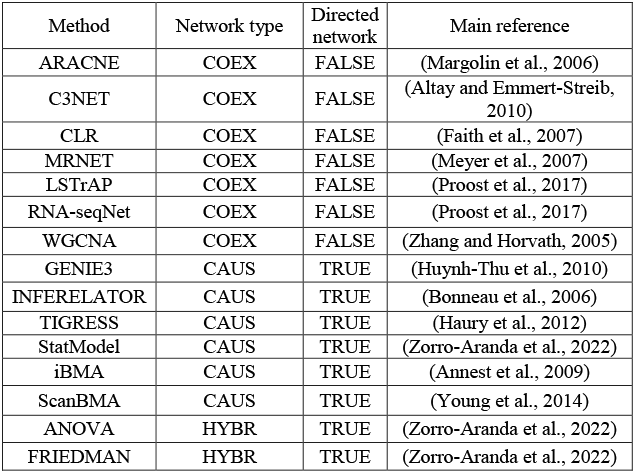
GRNs inference tools used in this work. For a detailed description of each tool, please see the Supplementary Files. COEX tools infer undirected networks, CAUS tools infer directed networks, HYBR tools infer undirected networks and the direction TF-TG is assigned with the list of known regulators to keep only the TF-mediated interactions (Zorro-Aranda et al., 2022). Community is not listed here because rather than a stand-alone tool, this method integrates the interactions from several single-tool predictions.

### Tools for inferring co-expression networks should be assessed apart from those for inferring causality

We used synthetic and biological datasets to assess the tools inferring networks from microarray data (Figure 2A). We assessed the inferred networks using 30 synthetic gene expression datasets with varying noise levels and sample sizes against the biological regulatory network used to generate the synthetic data. There was an overall improvement with larger datasets with less noise (Figure 2B and Supplementary Figure 2). GENIE3 and Inferelator performed the best, even better than Community, contrasting with the results of the DREAM5 challenge where Community outperformed all the single-tool predictions on the assessment with synthetic data (Marbach et al., 2012). On the other hand, ANOVA and WGCNA showed poor performance despite the data variations. There was no clear difference among the tools at the group level.

**Figure 2.**
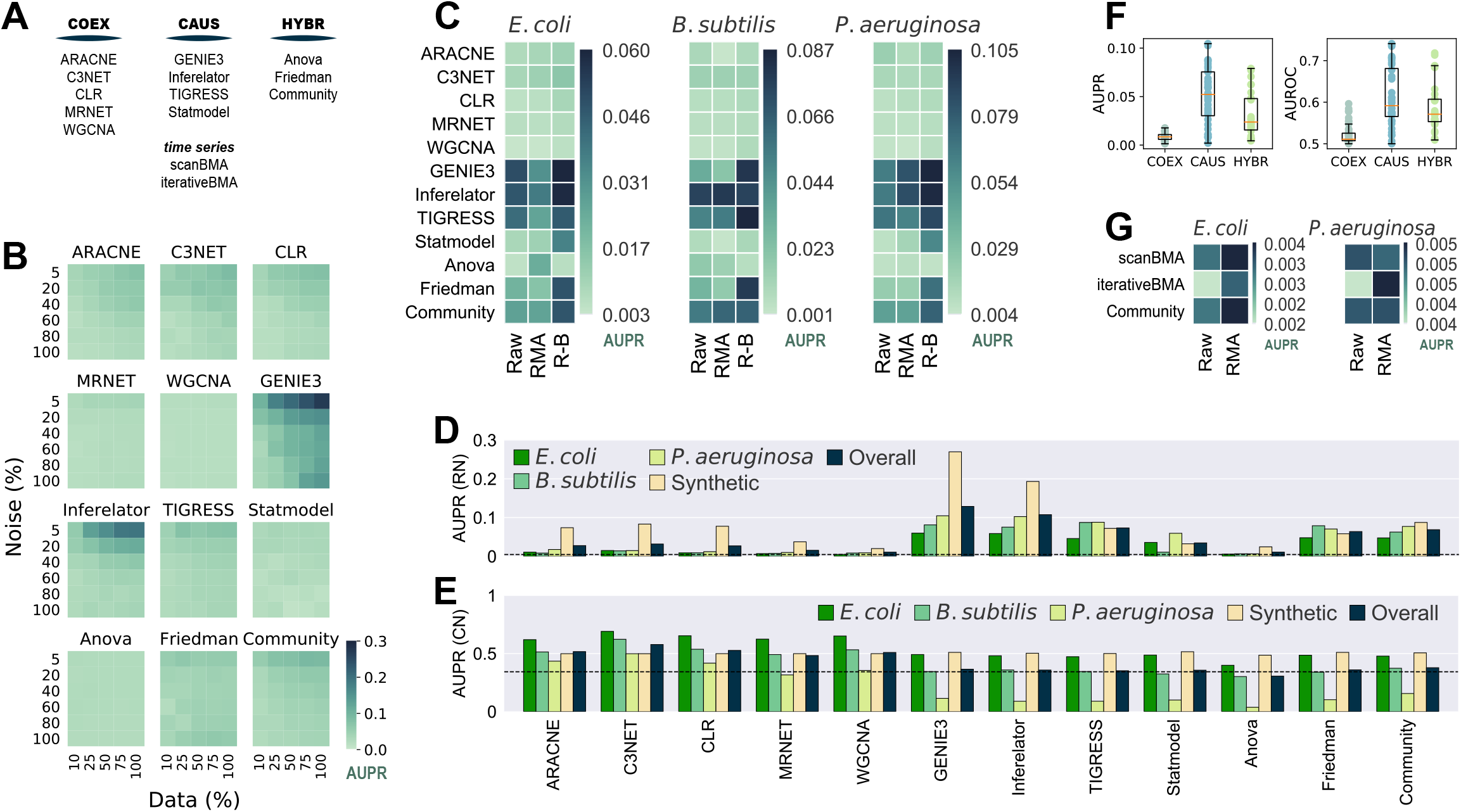
Assessment of network inference tools for microarray data. 100% of the synthetic dataset contains a total of 788 conditions. The Community Network is the integration of the single-tool predictions using the Borda count method (Marbach et al., 2012). **A)** Network classification. Network inference tools for microarray data were classified according to the type of network they infer. **B)** GENIE3 is the best tool for synthetic data. Synthetic gene expression datasets with different levels of noise and completeness were generated from the biological network of *E. coli* (511145_v2017_sRDB16_eStrong). The same network was used as the GS for the assessment. **C)** Batch correction and knowledge of the transcription factors improve the inference of transcriptional GRNs. Causal and Hybrid tools outperformed Co-expression tools in the assessment of GRNs using biological data for *E. coli*, *B. subtilis*, and *P. aeruginosa* with different levels of data normalization: raw data, Robust Multiarray Averaging (RMA), and RMA plus batch correction. Inferences were assessed with experimentally-validated GRNs. **D)** GENIE3 is the best tool for the inference of GRNs. **E)** Assessment for the inference of co-regulation network. The COEX tools outperformed CAUS and HYBR tools. C3NET performed the best. **F)** Boxplot representation of data in panel C to highlight the differences across tool groups. **G)** scanBMA outperformed iterativeBMA with biological data. The Community network for this panel only integrates interactions from scanBMA and iterativeBMA.

We collected gene expression data for *E. coli*, *B. subtilis*, and *P. aeruginosa* from GEO and generated three datasets for each organism, each with different preprocessing levels: raw data, Robust Multiarray Averaging (RMA) normalization, and RMA normalization plus batch correction (R-B). For the GS, we retrieved experimentally-supported GRNs from Abasy Atlas for the three organisms. As a group, CAUS performed the best followed by HYBR. On the other hand, COEXP showed poor results. Among the CAUS tools, GENIE3, Inferelator, and TIGRESS performed the best across the three organisms. GENIE3 was the best method in *E. coli* and *P. aeruginosa*, but TIGRESS and Inferelator outperformed it in *B. subtilis*, the organism with the smallest dataset (Supplementary Figure 3). This could be due to the lower prediction stability of GENIE3 to data size variations in contrast with TIGRESS and Inferelator. Among the HYBR tools, Friedman and Community improved their performance with R-B data, while ANOVA showed inconsistent results. Most of the tools performed better with fully preprocessed R-B (Figure 2C).

For each inference tool, we averaged its prediction score with the highest-quality data: R-B for each organism and the complete synthetic dataset with the lower noise level (5%). GENIE3 obtained the highest overall score, followed by Inferelator and TIGRESS (Figure 2D). Community ranked fourth in the overall score despite it includes interactions from the COEX predictions. In concordance with the DREAM5 challenge (Marbach et al., 2012), this suggests that despite low-scored predictions integration, Community still has reliable performance. A community integration seems to be a safer choice because the rank of individual tools differs among organisms, but CAUS tools outperformed COEX tools with biological data every time (Figure 2D).

Unlike the COEX tools, the CAUS and the HYBR tools require a list of the genes coding for TFs (Table 1) to keep only TF-TG interactions and avoid TG–TG edges that are not expected in a GRN, such as the networks used as GS. On the other hand, only a few of the interactions inferred by the COEX tools include a TF, i.e., most edges are TG-TG interactions (Supplementary figure 4). As an effort to perform a fair assessment of COEX tools, we modified the *E. coli* GS to resemble a co-regulation network where each regulon, set of co-regulated genes, is a clique (every node is interconnected). This way, COEXP outperformed the rest of the tools (Figure 2E).

The performance of every tool declined with the biological datasets in contrast to the synthetic ones. It is expected because the synthetic datasets were generated with the network used as GS. Besides, training and evaluating the tools with biological data is rare due to data accessibility (Marbach et al., 2010). There is a clear difference between the performance of CAUS and COEX tools with the biological datasets and a GRN as the GS (Figure 2C, F). On the other hand, the COEX tools succeeded with a simulated co-regulation network as the GS (Figure 2E). C3NET obtained the highest overall score, followed by CLR, ARACNE, and WGCNA. These results suggest that even though we should use CAUS tools for the inference or GRNs, tools inferring co-expression networks should be assessed apart from those inferring causality. Ignoring the direction of the GS interactions to make a fairer comparison (Chen and Mar, 2018) is not enough. Because of the nature of the network, the interactions inferred by COEX tools will be closer to representing co-expression and co-regulation rather than regulation. Moving to regulation is not trivial, but some approaches are already trying to infer causality from co-regulation and co-expression networks (Aibar et al., 2017; Chen and Liu, 2022).

Inference methods based on Bayesian approaches take advantage of time-series data to infer causal relationships (Lo et al., 2012). We assessed two tools based on a Bayesian approach: scanBMA (Young et al., 2014) and iterativeBMA (Annest et al., 2009), along with a Community reconstruction integrating both predictions. The performance with synthetic data improved with larger datasets and less-noise levels. iterativeBMA obtained the best scores, slightly better than Community (Supplementary Figure 5). Then, we assessed the tools with biological data, one time-series experiment for *E. coli* and one for *P. aeruginosa*. We used only raw (non-normalized) and RMA pre-processing steps as batch correction is not necessary for the one-source samples. Overall, scanBMA performed better than iterativeBMA (Figure 2G). Both tools with Bayesian approaches performed poorly despite their advantage over other methods to infer causal relationships, perhaps because of the few samples available. Future data availability along with experimental annotation might improve the performance of Bayesian approaches.

### RMA with Batch correction on large datasets improves the predictions

To provide deeper insights into the effects of data normalization on network inference, we contrasted the results using none (raw), RMA, and R-B preprocessing levels. The removal of batch-effect over RMA (R-B) normalization seems to slightly improve the predictions (Figure 3A and Supplementary figure 6). RMA normalization without batch correction worsens the performance of the tools. This is because some tools might be leveraging data heterogeneity or information lost in the normalization process (Sirbu et al., 2010). Besides, the assumptions considered by normalization pipelines could be violated, resulting in spurious predictions (Evans et al., 2018). Therefore, either raw data or normalized and batch-effect-corrected data should be used for network inference with highly heterogeneous datasets.

**Figure 3.**
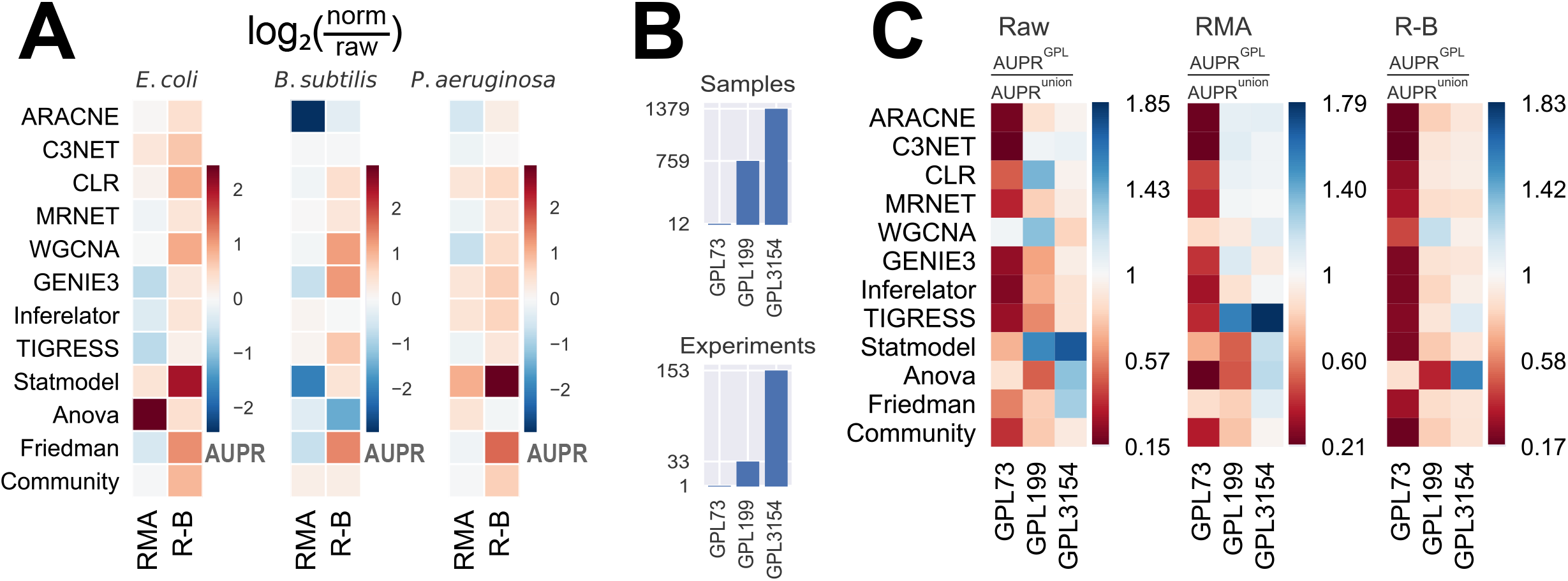
Effect of normalization and batch correction on the GRN inference with biological data. **A)** RMA normalization with batch correction (R-B) presents a slight improvement over only RMA normalization. The values represent the log2 ratio of the AUPR with normalized data concerning the AUPR with raw data. Higher (warmer) values mean more significant improvement with normalization. **B)** Platforms vary in the number of samples and experiments. **C)** Methods were assessed using different Affymetrix platforms of *E. coli* and. AUPR increases with larger datasets as input data.

In addition to data preprocessing, the dataset size should be considered a relevant factor in the prediction outcome. The dataset for *E. coli* was collected from three GEO platforms with a different number of samples (see Materials and methods): GPL73 (12 GSM), GPL199 (759 GSM), and GPL3154 (1379 GSM) (Figure 3B and Supplementary figure 7). We assessed the predictions using individual GEO platforms with the three preprocessing levels as input (Figure 3C and Supplementary figure 8). In general, there is an improvement in the prediction scores for larger datasets. The scores with GPL199 and GPL3154 are considerably higher than the score for the smallest platform (GPL73). However, there is not a remarkable difference between GPL199 and GPL3154 with RMA and R-B normalization. In the case of raw data, it seems to be an improvement as the data size increases. From these results, we can conclude that the larger the dataset the better the predictions. However, previous studies have shown that not only the dataset size but also the variability of conditions are relevant factors for network inference (Sastry et al., 2019). This is evident with the smallest platform which seems to have less heterogeneity among the platforms. In contrast, the other two platforms have better results alone than together which suggests that both have redundant information. Otherwise, normalized datasets with a size of two orders of magnitude would be good enough for network inference. These results are consistent across the three tool groups.

### A network-type-driven selective Community is the best choice when a GS is not available

A previous DREAM challenge suggested that integrating multiple single-tool predictions into a community network is a safe choice, especially when there is no partial network to use as GS (Marbach et al., 2012). Even though the AUPR and AUROC tend to be constrained to higher values as more single-tool predictions are integrated (Supplementary figure 9), the probability of CAUS tools outperforming Community decreases when their predictions are merged with other single-tool predictions (Figure 4A and Supplementary figure 10). This is due to the poor predictive power of some tools, which perform better only when integrated with several other predictions (e.g., ANOVA). The beginning of the prediction list is critical for the performance of the tools (Marbach et al., 2010). While COEX tools tend to have their true positive interactions scattered throughout the entire prediction, CAUS tools include most of their true positive interactions from the beginning (Supplementary figure 11).

**Figure 4.**
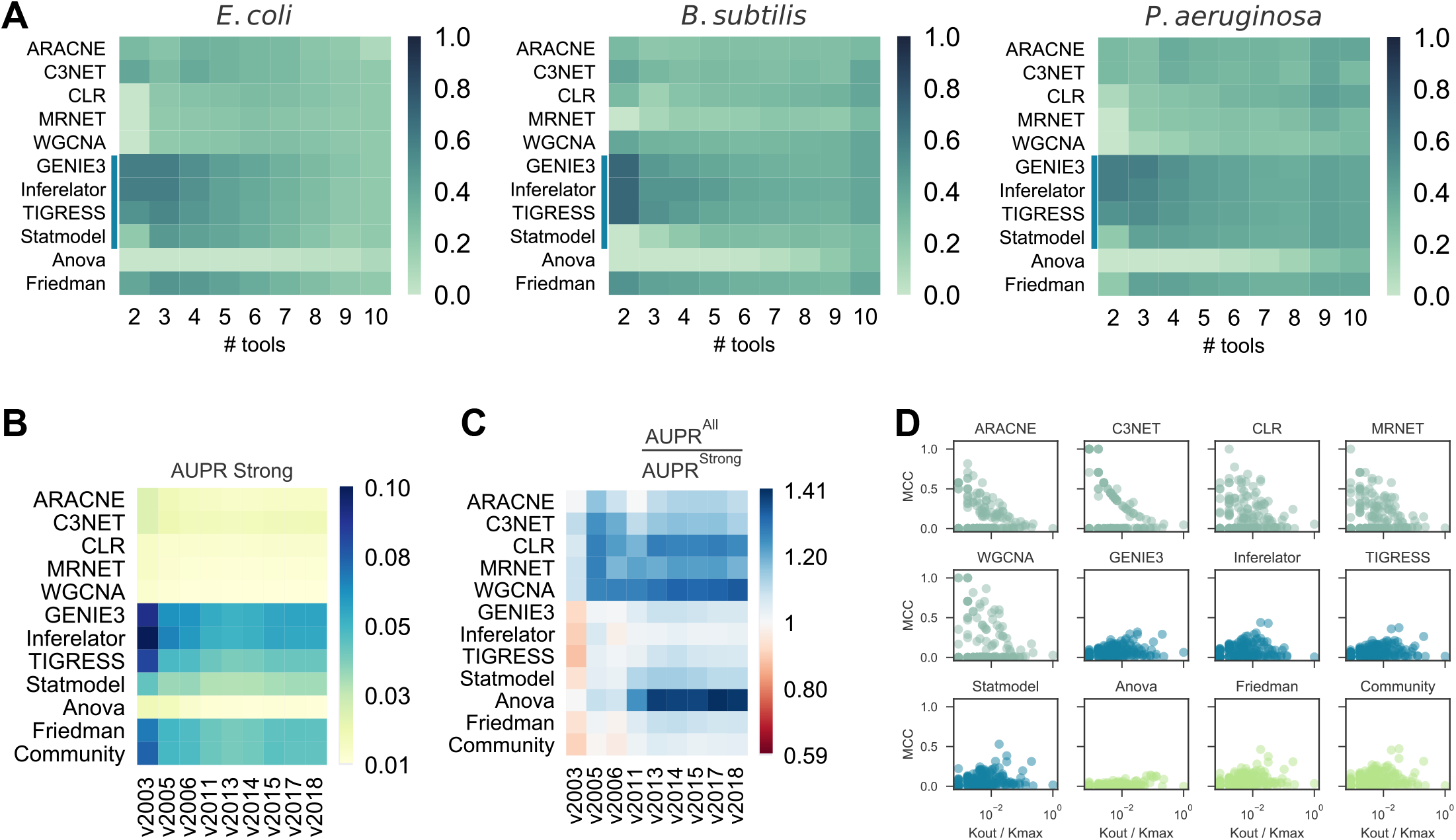
Effects of results integration, GS incompleteness, and Regulon-level assessment. **A)** Probability of a tool to outperform Community by its integration with others (# tools) into a selective community. CAUS tools are affected rather than improved by others. **B)** Assessment of GRN inference methods with the historical reconstruction of the *E. coli* GRN. The incompleteness of the GRN used as GS does not affect the AUPR score. **C)** AUPR ratio between a “strong” GS and a “weak” one. In most cases, the tools performed better when a “weak” GS was used. The “weak” GS is a superset of the “strong” GS including interactions supported by non-directed experiments. **D)** Regulon prediction assessment with Matthew’s correlation coefficient (MCC). Each dot represents a regulon inference for an *E. coli* TF, higher is better. Out-degree connectivity (Kout) for the TF controlling the regulon is normalized by the maximum connectivity (Kmax) of the *E. coli* network.

### COEX tools capture function-specific regulons and non-direct interactions

We assessed the predictions with snapshots of the historical reconstruction of the *E. coli* GS, each of these networks with two versions; one with all the interactions discovered at a specific timepoint (“all”) and the other one with only validated protein-DNA interactions (“strong”). The assessment methodology showed robustness to the incompleteness of the GS (Figure 4B), suggesting that CAUS tools outperform COEX tools with every snapshot of the GS, disregarding its completeness level. Moreover, even though all the tools improved the performance with the “all” GS, the difference is bigger for COEXP tools (Figure 4C). While the “strong” GS only contains direct TF-DNA interactions, the “all” GS may contain non-direct interactions (i.e., an interaction mediated by a third biological entity) (Escorcia-Rodriguez et al., 2020). Gene expression data capture both direct and non-direct regulatory events. Therefore, inference tools based solely on gene expression data tend to also infer non-direct interactions, especially COEX tools (Figure 4C). Perhaps, this consideration may shed light on the search for consistency between GRNs and gene expression data (Larsen et al., 2019; Parise et al., 2021). On the other hand, every tool performs better with the “strong” GS on AUROC (Supplementary figure 12), but this is because of the highly unbalanced positives/negatives ratio (Saito and Rehmsmeier, 2015).

We assessed the predictions at the regulon level using the F1 score. The CAUS tools performed better on large regulons (i.e., those of global regulators) (Supplementary figure 13). On the other hand, the COEX tools are the best alternative for local regulators, which are associated with function-specific regulons (Freyre-Gonzalez et al., 2022). To discard potential bias induced by the F1 metric (Chicco and Jurman, 2020), we also used Matthew’s correlation coefficient (MCC), obtaining consistent but less meaningful patterns (Figure 4D). The explanation for this is that COEX tools distribute the interactions among all the genes sub-estimating the number of TGs for global regulators, while CAUS and HYBR tools distribute the interactions only among the TFs list provided over-estimating the number of TGs for each TFs, especially for local TFs (Supplementary figure 14).

### Unsupervised learning with global structural properties segregates COEX inferences from the rest of the networks

Beyond assessing the tools solely based on the standard statistical metrics, we analyzed global structural differences among the networks. We computed the following structural properties for the regulatory networks: density, number of regulators, maximum out-connectivity, feedforward and complex feedforward circuits (Alon, 2007; Freyre-Gonzalez and Tauch, 2017), 3-feedback loops, size of the giant component, average clustering coefficient, diameter, average shortest path length, and the coefficient of determination for the degree P(k) and clustering coefficient distribution C(k) (Albert, 2005). See Supplementary file 3 for the definition of the structural properties. Then, we clustered the networks based on their normalized global structural properties (Materials and methods).

For the *E. coli* networks, COEX tools were clustered into one group (Figure 5A). On the other hand, CAUS and HYBR tools were clustered into a second group, excluding ANOVA. Even though the GS was not clustered with any of the two major groups, it was closer to the latter one (Figure 5A). We obtained similar results with the networks for *B. subtilis* (Figure 5B) and *P. aeruginosa* (Figure 5C), although the GS for *B. subtilis* got much closer to the CAUS and HYBR group (Figure 5B).

**Figure 5.**
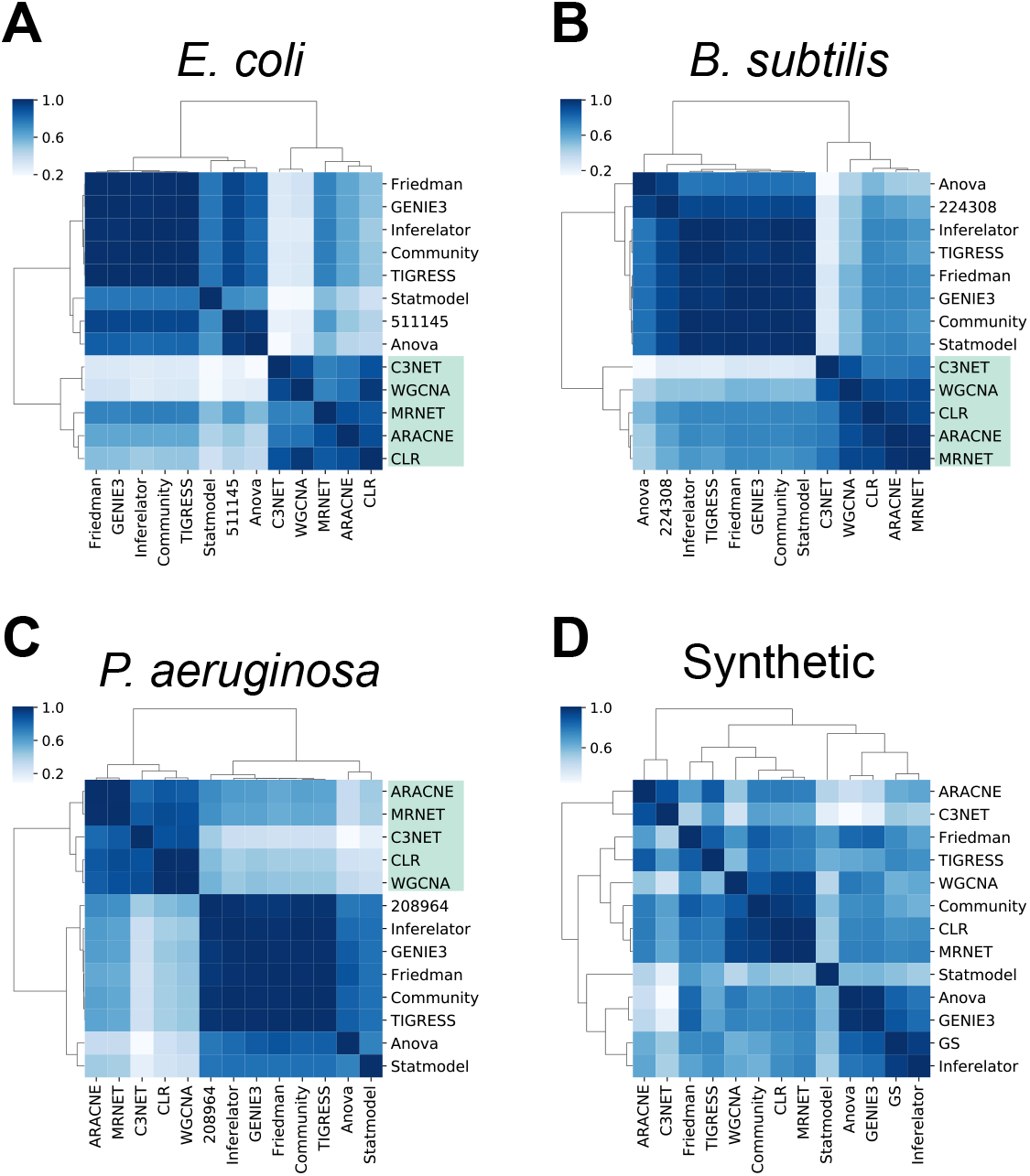
Assessment of the structural properties. Clustering of the global structural properties suggests there is a clear structural difference between causal (CAUS) and non-causal (COEX) networks. In contrast to the inferences with biological data (A-C), most networks inferred from synthetic data (D) are more similar to each other.

The clusters were not conserved with synthetic data inferences, suggesting that inferences with synthetic data were structurally similar disregarding their type of interactions (Figure 5D). Contrary to biological data, GeneNetWeaver generates the synthetic data following the topology of the network provided (Schaffter et al., 2011), making it easier for the tools to recover such topology. Several structural properties are constrained by the graph complexity and characterize the GRNs with causal interactions, despite different network completeness levels (Campos and Freyre-Gonzalez, 2019; Escorcia-Rodriguez et al., 2021). Therefore, we expect such properties to remain similar in the final GS, and the overall topological assessment of the predicted networks will be like the one performed with the current GS.

We then used an in-house Python implementation of the previously reported *D*-value (Schieber et al., 2017), which assesses network similarity based on topological evidence taking centrality into account. For the biological datasets, CAUS tools were always clustered with Community and Friedman but never with the GS (Supplementary Figure 15). Noteworthy, the GS was not clustered with the COEX tools either. Instead, it was isolated, as well as the ANOVA network. Overall, the results remain consistent across organisms, clustering CAUS networks apart from the COEX ones. Further topological analysis with all historical GS for *E. coli* showed that, despite GS completeness, the same conclusions are expected (Supplementary figure 16).

Noteworthy, two networks might have identical global structures with no intersection between their regulations (shuffled node labels). This explains why ANOVA was repeatedly clustered with the GS, despite its poor performance with standard assessment metrics. However, between the two strategies to assess the structure of the networks, the one based on the normalized structural properties in GRNs (Figure 5) is more consistent with the standard metrics. We suggest using this approach as a complementary assessment always a GS is available. When no GS exists for the organism of interest, the structural assessment can be used along with other biological networks and random models to prove the prediction is structurally more similar to a biological network than random. We recently used this approach to assess network inferences for *Rhizobium etli* (Taboada-Castro et al., 2022).

Analyzing the structural properties individually (**Table 2** and Supplementary figure 17), COEX tools have higher density and fraction of regulators. Given that the predictions have the same number of interactions, having a higher fraction of regulators results in lower max out-connectivity. On the other hand, synthetic predictions tend to have higher max out-connectivity values than their biological counterparts. Noteworthy, the max out-connectivity for the *P. aeruginosa* GS might be underestimated due to low genomic coverage (Escorcia-Rodriguez et al., 2020). Regarding normalized path-related properties, the COEX tools have the largest normalized diameter and average path length due to their small fraction of nodes in their giant component (Table 2). Contrary to COEX tools that reach more than 200 components, CAUS and HYBR tools predict networks with a few components (Supplementary figure 18) because their maximum is constrained to the number of TF-coding genes; and every interaction connecting regulons decreases the number of components. A high P(k) coefficient of determination (R^2^) value was found across all biological predictions and all GSs. The C(k) R^2^ was good only for COEX and HYBR biological predictions suggesting their modularity, like the one found in the GS. Regarding network motifs, the COEX inferences were the most similar to the GS. This agrees with the motifs search in DREAM5 where feed-forward loops were recovered most reliably by mutual-information and correlation-based methods (Marbach et al., 2012) (i.e., COEX tools).

### GENIE3 outperformed tools developed for bulk RNA-seq

We interrogated the performance dependence of GRNs inference related to transcriptomic technique, comparing two COEX inference tools developed exclusively for bulk RNA-seq data (RNAseqNet (Imbert et al., 2018) and LSTrAP (Proost et al., 2017)) and the best CAUS microarray-based approach (GENIE3). We retrieved RNA-seq datasets for *E. coli* and performed a cross-evaluation between the tools, exchanging the input data. First, we used a subset (see Materials and methods) of our raw and RMA microarray datasets of *E. coli* to reduce the impact of data size variation and observed that GENIE3 outperformed RNASeqNet and LSTrAP significantly (Supplementary figure 19). Next, we used the RNA-seq datasets (raw counts, normalized with DESeq2, and PRECISE (Sastry et al., 2019)) as input. The COEX RNA-seq-based tools performed better with the homogenous largest RNA-seq dataset, PRECISE (Supplementary figure 19). Despite the improvement of RNASeqNet and LSTrAP with the RNA-seq data, GENIE3 still performed better (Supplementary figure 19). These results agree with a previous synthetic gold standard-based benchmarking of network inference methods for scRNA-seq data where GENIE3 is still placed within the top-performing tools methods (Pratapa et al., 2020), making GENIE3 a top-performing tool regardless of the transcriptomic technique.

Furthermore, we assess the predictions based on their global structure (Supplementary figure 20). We only considered the inferences datasets with the best MCC scores (Supplementary figure 19), PRECISE for RNA-seq, and raw for microarray data. Both datasets and metrics showed consistent results clustering the GS with GENIE3, RNAseqNet, and Community, leaving LSTrAP out (Supplementary figure 20). This suggests that RNAseqNet infers networks with structural properties more similar to the GS than LSTrAP does. However, non-ranked interactions might be a shortcoming for RNAseqNet.

Overall, compared to how well the tools performed with microarray data, RNA-seq data did not significantly improve their performance. It agrees with a previous assessment in *A. Thaliana*, where networks derived from simple correlations and microarray data obtained higher scores than inferences with RNA-seq data (Giorgi et al., 2013). Although RNA-seq has progressively replaced microarrays (Lowe et al., 2017), the gene coverage referred to as an advantage of RNA-seq, is less of a problem for microarrays in model prokaryotes where new microarrays have overcome the coverage issue (Swarbreck et al., 2008). Despite the amount of available RNA-seq data, most organisms do not have an appropriate annotation (Salzberg, 2019), while large microarray-based transcriptomic data have continuously grown into public databases (Barrett et al., 2013; Athar et al., 2019).

### Conclusions and guidelines

All the CAUS tools (GENIE3, TIGRESS, Inferelator, and Statmodel) outperformed the COEX tools when assessed with a GRN as the GS (TF–TG interactions) with biological and synthetic data and, taking advantage of a TFs list. Even though we filtered TF-TG interactions from the co-expression inferences approaches (HYBR), the performance of CAUS tools was still better. GENIE3 and Inferelator performed the best for synthetic and biological data. GENIE3 also outperformed inference tools developed for bulk RNA-seq data. COEX tools performed better when assessed with a GS resembling a co-regulation network (interactions among co-regulated genes). Regarding time-series tools, scanBMA performed the best, although it is highly affected by dataset size.

Larger datasets result in better predictions but require a selective inferences-integration process and batch correction to mitigate technical variability; applying only RMA worsened the predictions. The probability of CAUS tools outperforming Community decreases as more tools are integrated into a community network, suggesting the use of a selective community based on the desired output network type (co-regulation or GRN). Although CAUS tools are the best alternative to infer global GRNs, COEX tools are better at inferring regulons for function-specific (i.e., local) TFs. An assessment against a GS including potential indirect interactions suggests that COEX tools might be the best alternative to identify indirect regulations. This is useful when the goal is to identify all the regulators affecting the expression of a gene, and not only DNA-binding TFs (Zorro-Aranda et al., 2022).

Based on global structural properties, COEX tools segregate from CAUS tools when using biological predictions, highlighting the differences among their output network type. Individual structural properties support the similarity between CAUS inferences and the GRNs used as GS. However, no clear clusters were found with synthetic data, suggesting that biological data is required for the structural assessment because synthetic data generation is based on the topology of the input network (Schaffter et al., 2011). Historical snapshots of the GS suggest the statistical and structural assessment to be robust to GS incompleteness.

The overall modest performance of the tools is evident and the potential inherent pitfalls to the conjecture that statistical relationships between expression profiles correspond to regulatory interactions have been previously noted (Pratapa et al., 2020; Freyre-Gonzalez et al., 2022). Recent works leveraging prior networks, structural constraints, and sequence motifs to improve transcriptomic-based GRN inference have shown promising results (Castro et al., 2019; Lim et al., 2022; Zorro-Aranda et al., 2022). Following we provide guidelines for the inference and assessment:

### Inference

- Identify the best kind of tool for your purposes.
  - CAUS or Community for whole GRNs or regulons of global TFs.
  - COEX for regulons of local TFs (few targets), co-expression, or co-regulation networks.
  - Using a list of TFs to filter inferences based on co-expression (e.g., ANOVA and Friedman) to get a causal network is not enough to infer a global GRN. Integrate the TFs into the inference pipeline.
- A selective community based on the type of network required is better than an all-inclusive community.
- If you want to use one COEX tool, use C3NET but keep in mind you will obtain a co-expression network, not a GRN.
- If you want to use one CAUS tool, use GENIE3 disregarding the type of gene expression data used.
- Merge datasets only when the final size of the data outweighs the noise of merging different sources.
- In prokaryotes, dataset size and preprocessing are more important than the transcriptomic technique used to generate the data.
- Normalize your data using Batch correction if it is necessary. Using only RMA is not recommended.
- If it is feasible, take advantage of biological information such as a list of TFs.

### Assessment

- Using synthetic data to assess the predictions might provide insights about the performance of the tools but expect it to worsen when assessed with biological data and the inferred networks to have a different global structure.
- Take advantage of several experimentally-validated bacterial GRNs to be used as GS (e.g., https://abasy.ccg.unam.mx/ for bacteria).
- Whenever possible, use historical snapshots or network sampling to prove the results hold despite GRN incompleteness.
- Use MCC to perform a regulon-level assessment of the predictions.
- Compare network structural properties to assess the global topology of the networks inferred from biological data.
- A structural assessment of the predictions applies to biological data only. Because of the mechanisms to generate the data following the topology of an input network, predictions with synthetic data have a similar structure despite inherent differences.

## Materials and methods

### Selection of GRN prediction methods to be assessed

We thoroughly reviewed the literature and selected methods that were able to infer a GRN from an expression data matrix. We also considered usability, which takes into account 1) open-source availability, 2) enough documentation, and 3) the ability to be run by a command line.

### Synthetic data

The synthetic datasets, all with 788 conditions (rows) and 197 genes (columns), were generated using GeneNetWeaver software (Schaffter et al., 2011) applying the standard procedure reported by the DREAM5 consortium, with the *E. coli* network (511145_v2017_sRDB16_eStrong) from Abasy Atlas (Escorcia-Rodriguez et al., 2020) being used as the seed. To explore the effects of noise levels in GRN inference, we generated datasets with 20%-step values for the noise parameter, as well as the 5% noise level selected for the DREAM5 challenge. To study the effect of sample size in GRN inference, we sampled each of the previous datasets at 10, 25, 50, 75, and 100% of experimental conditions, preserving an equal representation of each experimental condition. The same procedure was followed for time-series 4207 conditions and 197 genes data generation.

### Data extraction, normalization, and batch-effect correction for biological data

#### Microarray data

The microarray data for *Escherichia coli* K-12 MG1655, *Bacillus subtilis subtilis* 168, and the pathogen *Pseudomonas aeruginosa* PAO1 were retrieved from the (GEO) database using four main inclusion criteria: A) records were associated with public Affymetrix platforms and had an available CEL file useful to perform Robust Multi-chip Averaging (RMA) normalization by Oligo R package (array annotation package); B) an available GEO Series Matrix, an expression matrix annotated as non-normalized data, referred here as raw data, and C) more than one available sample. In addition, we excluded GEO samples related to more than one organism. For *E. coli*, a total of 2,154 GEO samples (GSM) from 182 GEO Series (GSE) were retrieved from the GEO platforms GPL73 (1 GSE, 12 GSM), GPL199 (33 GSE, 759 GSM), and GPL3154 (153 GSE, 1379 GSM). After applying RMA, we kept with the shared genes among *E. coli* GPLs belonging to the K-12 MG1655 strain, obtaining a total of 4003 genes, which comprise 87.7% of the genome. For *B. subtilis* we used the platform GPL343 and retrieved 7 GSE with a total of 64 GSM, obtaining a total of 4010 genes, which comprises 88.5% of the genome. Finally, for *P. aeruginosa* we used the GPL84 platform with 125 GSE and a total of 1133 GSM, obtaining a total of 5548 genes, which comprise de 97.4% of the genome. Microarray raw data (CEL files) were normalized through the RMA implementation in the R package *oligo* (Carvalho and Irizarry, 2010), using default parameters. Next, we removed all the conditions in which NANs or zeros were present due to normalization effects. Lastly, we performed a batch-effect correction using ComBat (Johnson et al., 2007) implementation in the sva R package with a non-parametric adjustments approach (function from the sva R package using the following parameters: mod = NULL, par.prior = FALSE, mean.only = FALSE).

#### Time-series microarray data and condition sampling

Since GEO does not provide a feasible way to filter TS experiments, we used all public metadata of samples to identify GSE records with a timeline progression and filtered them with our inclusion criteria. From the identified TS GSE list we selected the largest record for each organism. For *E. coli we* used GSE12411 and retrieved 28 GSM with 3 time-points with 4, 12, and 12 replicates respectively, regarding *P. aeruginosa* we used GSE52445 with 28 GSM representing 14 time points each one with 2 replicates. For the assessment, we used only raw and RMA preprocessed data, the batch correction step was not necessary for the one-source samples.

We sampled the Abasy Atlas networks (Escorcia-Rodriguez et al., 2020) to allow dimensionality reduction by the Bayesian tools (Annest et al., 2009; Young et al., 2014). We sampled the networks 511145_v2018_sRDB18_eStrong for *E. coli* and 208964_v2015_s11-RTB13 for *P. aeruginosa*. We applied snowball sampling (Heckathorn and Cameron, 2017), also known as link-tracing, using the network nodes with the highest degree of centrality as seed and 198 as the cutoff value for the sampling to get the same size of data as in the *in silico* time-series assessment. The final sample sizes were 139 samples x 198 genes for *E. coli* and 45 samples x 198 genes for *P. aeruginosa*. We used 198 genes for consistency with the time series synthetic data.

#### Methods, data collection, and assessment for cross-evaluation

To compare the performance dependency of the RNA-seq-based and microarray-based inference methods, we swap their transcriptomic input data and compare it with the original correspondence input results. Due to the diversity of RNA-seq-based methods, we preselected LSTrAP, RNAseqNet, and VCNet exclusively developed for GRN inference from bulk RNA-seq. However, we excluded VCNet from the analysis since it cannot be applied to a large number of samples unless you optimize the computational complexity inherent in its loop-based code. On the other hand, RNAseqNet and LsTrAP are low-time-consuming algorithms that aim to increase the reliability of inference from biologically related datasets (Imbert et al., 2018).

#### Bulk RNA-seq data

We collected two bulk RNA-seq datasets for *E. coli* K-12 MG1655. The small one was retrieved from GEO NCBI (GSE73673) (Kim et al., 2016), we downloaded the 87 sample files with the processed reads (Kim et al., 2016) for 3923 genes. Next, we applied the DESeq2 normalization (Love et al., 2014) a commonly used method that has been evaluated against different normalization methods (Dillies et al., 2013; Maza et al., 2013; Soneson and Delorenzi, 2013; Smid et al., 2018). For the largest one, we download the available processed (log_tpm.tsv) dataset from PRECISE 1.0 (Sastry et al., 2019), a Precision RNA-seq Expression Compendium for Independent Signal Exploration, build it with 15 studies derived with a standardized protocol from the same research group and PRECISE developer. We only kept with the genes shared between the PRECISE dataset and our microarray dataset resulting in 278 conditions and 3557 genes.

#### Microarray data transformation

We sampled a subset of 87 samples from our collected *E. coli* microarray dataset. We used only the raw and RMA version since batch correction was not applicable. Unfortunately, the RNAseqNet algorithm takes as input read counts or TMM normalized counts data; thus, we avoided negative values from sampling. To the best of our knowledge, RNAseqNet is not able to work with microarray or RNA-set datasets without filter genes with at least 70% of sample coverage.

#### Gold standards

We used strongly-supported, meta-curated GRNs from Abasy Atlas v2.2 (Escorcia-Rodriguez et al., 2020) as GSs for *E. coli* (511145_v2018_sRDB18_eStrong), *B. subtilis* (224308_v2008_sDBTBS08_eStrong) and *P. aeruginosa* (208964_v2015_s11-RTB13). The nodes of Abasy GRNs depict either genes, regulatory sRNAs, or regulatory protein complexes. For this work, we converted networks with genes and regulatory protein complexes into gene-gene networks to use as GS since only those interactions can be inferred. We removed the genes for which no expression data was retrieved since the prediction of its interactions would not be inferred by the methods assessed in this work. We obtained a total of 4075 interactions among 1780 genes for *E. coli*, 2294 interactions among 1298 genes for *B. subtilis*, and 1297 interactions among 868 genes for *P. aeruginosa*. For GS incompleteness analysis, we also retrieved from Abasy various public versions of the *E. coli* GRN (hereafter referred to as historical snapshots), with different completeness levels.

For the construction of the GS with interactions between co-regulated genes, we connected each regulon of 511145_v2018_sRDB18_eStrong so each of them forms a clique and obtained a total of 737913 interactions between the same number of genes, overestimating the density of the network. Note that, in such network representation, the TGs from a regulon formed a clique, including the TF only if it regulates its own transcription. For the synthetic GS, we used 511145_v2017_sRDB16_eStrong as input for GeneNetWeaver (Schaffter et al., 2011) to generate datasets with 5,20,40,60,80, and 100% noise variations. From such datasets, we generated subsamples with 20,25,50,75, and 100% completeness.

#### Integration of individual predictions into a Community network

A confidence score provided by each tool (when available) was used to rank predictions and missing interactions were ranked right after the last predicted one. Therefore, longer predictions penalize more the missing interactions. Inferred interactions sharing a common score by a method were ranked equally. The average rank is used as a score for the Community. For biological data, predictions were previously trimmed to the number of interactions that the complete organism-specific GRN would have according to previous work (Campos and Freyre-Gonzalez, 2019; Escorcia-Rodriguez et al., 2020). Those values correspond to 12000 for *E. coli* and *B. subtilis* and 16000 for *P. aeruginosa*.

#### Assessment

Unless otherwise described in the analysis, network predictions larger than the expected number of interactions in the complete GRN were trimmed (Campos and Freyre-Gonzalez, 2019; Escorcia-Rodriguez et al., 2020). The first 12000 inferred interactions were considered for the assessment with *E. coli* and *B. subtilis* and the first 16000 inferred interactions for *P. aeruginosa*. Interactions shaping the GS were used as the positive set (P), while interactions absent in the GS were labeled as the negative set (N). Inferred interactions were considered True Positive (TP) if they were present in the GS and False positive (FP) if otherwise. Interactions in the GS that were not recovered by the algorithm were considered False Negative (FN). The Area Under Receiver Operating Characteristics (AUROC) and Area Under Precision-Recall (AUPR) curves were used to assess the predictions. While AUROC represents the specificity (FP/N) and the sensitivity (TP/P) of the prediction compared with the whole set of potential interactions, AUPR focuses on the list of predictions and its precision (TP/(TP+FP)) as well as the sensitivity of the algorithm. We select PR as the main assessment measure, due to the imbalance between positive and negative sets (Saito and Rehmsmeier, 2015). For the overall score, we used the average score for the complete dataset with 5% of noise for the synthetic data and scores obtained with RMA plus batch effect correction for biological data. For the study of the effect of GS incompleteness, we used each historical snapshot of the *E. coli* GRNs as the GS. Inferred interactions sharing the same score were considered as equally ranked by the method and genes present neither in the GS nor in the expression data were not considered for this assessment. For the assessment of predictions not providing a score for each interaction, we used the MCC which is the best-suited coefficient for imbalanced datasets (Boughorbel et al., 2017). Note that MCC was used only for the comparative assessment between GENIE3 and RNAseqNet, and the regulon-level assessment. RNAseqNet does not score the predictions. Therefore, we considered the first 12000 interactions to assess its prediction with MCC so the ranking of the interactions does not impact the score. Note that this is not ideal as the true positives –as well as novel interactions– may be at the bottom of the prediction making it disadvantageous for the experimental validation of such inferred interactions. For the regulon-level assessment, we trimmed the predictions to the expected number of interactions once the corresponding network is completed and compared each of the regulons against the cognate regulon in the GS using MCC and F1 score. The scores were plotting against the normalized out-connectivity.

For the prediction of the COEX methods, we duplicated every interaction in the prediction list, changing the direction. This is because the outputs provide interactions between two genes with no direction (e.g., symmetric adjacency matrix). Given the nature of the assessment with a directed network as the GS, we considered every interaction to be in both directions. While this would increase the number of predictions, consideration of the direction is required. On the other hand, for the assessment of the predictions with a co-regulation GS, we did not consider the direction of the interactions. This way, the direction of the interactions predicted by a CAUS method, was not considered.

#### Combinatorial

We constructed selective communities with the possible combinations of the 12 methods used in the assessment with biological data. We use the dataset normalized with RMA and batch correction for the three organisms. To measure the effect of each method on the community network, we computed the dominance score defined as the probability of a selective community network with a given tool outperforming the all-inclusive community network:

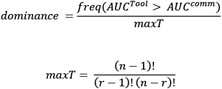

Where *maxT* is the theoretical maximum of selective communities with each tool, *n* is the number of methods (12) used for the combinatory, and r is the number of elements in the combinatory [2-11].

#### Structural properties

We computed several structural properties for GRNs at a global scale and normalized them as follows: Regulators, self-regulations, maximum out-connectivity, and giant component size were normalized by the network size (number of nodes). Density was used as its product with the fraction of nodes acting as regulators. Network diameter was normalized by the number of nodes – 2 (as if no shortcuts would exist). Network motifs were normalized by the number of potential motifs in the network, defined as:

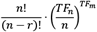

Where *n* is the size of the network, *r* is the number of nodes in the motif (*r* = 3), *TF_n_* is the number of TFs in the network, and *TF_m_* is the number of TFs required for each type of motif (*TF_m_* = 2 for feedforward and complex feedforward loops, and *TF_m_* = 3 for 3-feedback loops). We scaled the property values across networks between 0 and 1. We clustered networks and properties using Ward’s method. Further, we used pairwise Pearson correlation for the network property vectors and clustered them according to the Euclidean distance using Ward’s method.

We used an in-house Python implementation of the dissimilarity measure proposed by Schieber et al. (Schieber et al., 2017) to quantify the differences in the structural topology between two networks considering global structural properties, node-level structural properties, and centrality. We used the parameters the authors recommend (0.45,0.45,0.1) and applied used to compare the networks pairwise. The dissimilarity matrix was clustered using Pearson and ward’s method.

## Availability of supporting data

The data sets supporting the results of this article are available in the Abasy Atlas database (https://abasy.ccg.unam.mx/) and the supplementary material.

## Authors’ contributions

JE-R, Investigation, Methodology, Visualization, Writing—original draft, Writing—review and editing. EG-N Investigation, Visualization, Writing—original draft, Writing—review and editing. EH-B Investigation, Visualization, Writing—original draft, Writing—review and editing. AZ-A Investigation, Writing—original draft, Writing—review and editing. MT-P Investigation and Writing—original draft. JF-G Conceptualization, Funding acquisition, Investigation, Project administration, Resources, Supervision, Writing—review, and editing.

## Acknowledgments

We thank Roberto Olayo-Alarcón and Luis F. Altamirano-Pacheco for technical support in running some of the inference tools in a previous version of this work. JE-R is a doctoral student from Programa de Doctorado en Ciencias Biomédicas, Universidad Nacional Autónoma de México (UNAM). He received fellowship 959406 from CONACYT. A.Z.-A. received a doctoral scholarship [call 647 (2014)] from the Colombian Administrative Department of Science, Technology, and Innovation (COLCIENCIAS).

## Additional files

**Supplementary File 1.** Comparative table among the tools assessed in this work.

**Supplementary File 2.** A detailed description of the tools used in this work, supplementary figures, and references.

**Supplementary File 3.** Properties used for the structural assessment and their definition.

## References

Aibar, S., Gonzalez-Blas, C.B., Moerman, T., Huynh-Thu, V.A., Imrichova, H., Hulselmans, G., et al. (2017). SCENIC: single-cell regulatory network inference and clustering. Nat Methods 14(11), 1083–1086. doi: 10.1038/nmeth.4463.

Akesson, J., Lubovac-Pilav, Z., Magnusson, R., and Gustafsson, M. (2021). ComHub: Community predictions of hubs in gene regulatory networks. BMC Bioinformatics 22(1), 58. doi: 10.1186/s12859-021-03987-y.

Albert, R. (2005). Scale-free networks in cell biology. J Cell Sci 118(Pt 21), 4947–4957. doi: 10.1242/jcs.02714.

Alon, U. (2007). Network motifs: theory and experimental approaches. Nat Rev Genet 8(6), 450–461. doi: 10.1038/nrg2102.

Altay, G., and Emmert-Streib, F. (2010). Inferring the conservative causal core of gene regulatory networks. BMC Syst Biol 4, 132. doi: 10.1186/1752-0509-4-132.

Annest, A., Bumgarner, R.E., Raftery, A.E., and Yeung, K.Y. (2009). Iterative Bayesian Model Averaging: a method for the application of survival analysis to high-dimensional microarray data. BMC Bioinformatics 10, 72. doi: 10.1186/1471-2105-10-72.

Athar, A., Fullgrabe, A., George, N., Iqbal, H., Huerta, L., Ali, A., et al. (2019). ArrayExpress update - from bulk to single-cell expression data. Nucleic Acids Res 47(D1), D711–D715. doi: 10.1093/nar/gky964.

Barrett, T., Wilhite, S.E., Ledoux, P., Evangelista, C., Kim, I.F., Tomashevsky, M., et al. (2013). NCBI GEO: archive for functional genomics data sets--update. Nucleic Acids Res 41(Database issue), D991–995. doi: 10.1093/nar/gks1193.

Bellot, P., Olsen, C., Salembier, P., Oliveras-Verges, A., and Meyer, P.E. (2015). NetBenchmark: a bioconductor package for reproducible benchmarks of gene regulatory network inference. BMC Bioinformatics 16, 312. doi: 10.1186/s12859-015-0728-4.

Bonneau, R., Reiss, D.J., Shannon, P., Facciotti, M., Hood, L., Baliga, N.S., et al. (2006). The Inferelator: an algorithm for learning parsimonious regulatory networks from systems-biology data sets de novo. Genome Biol 7(5), R36. doi: 10.1186/gb-2006-7-5-r36.

Boughorbel, S., Jarray, F., and El-Anbari, M. (2017). Optimal classifier for imbalanced data using Matthews Correlation Coefficient metric. PLoS One 12(6), e0177678. doi: 10.1371/journal.pone.0177678.

Campos, A.I., and Freyre-Gonzalez, J.A. (2019). Evolutionary constraints on the complexity of genetic regulatory networks allow predictions of the total number of genetic interactions. Sci Rep 9(1), 3618. doi: 10.1038/s41598-019-39866-z.

Carvalho, B.S., and Irizarry, R.A. (2010). A framework for oligonucleotide microarray preprocessing. Bioinformatics 26(19), 2363–2367. doi: 10.1093/bioinformatics/btq431.

Castro, D.M., de Veaux, N.R., Miraldi, E.R., and Bonneau, R. (2019). Multi-study inference of regulatory networks for more accurate models of gene regulation. PLoS Comput Biol 15(1), e1006591. doi: 10.1371/journal.pcbi.1006591.

Chen, G., and Liu, Z.P. (2022). Inferring causal gene regulatory network via GreyNet: From dynamic grey association to causation. Front Bioeng Biotechnol 10, 954610. doi: 10.3389/fbioe.2022.954610.

Chen, S., and Mar, J.C. (2018). Evaluating methods of inferring gene regulatory networks highlights their lack of performance for single cell gene expression data. BMC Bioinformatics 19(1), 232. doi: 10.1186/s12859-018-2217-z.

Chicco, D., and Jurman, G. (2020). The advantages of the Matthews correlation coefficient (MCC) over F1 score and accuracy in binary classification evaluation. BMC Genomics 21(1), 6. doi: 10.1186/s12864-019-6413-7.

De Smet, R., and Marchal, K. (2010). Advantages and limitations of current network inference methods. Nat Rev Microbiol 8(10), 717–729. doi: 10.1038/nrmicro2419.

Dillies, M.A., Rau, A., Aubert, J., Hennequet-Antier, C., Jeanmougin, M., Servant, N., et al. (2013). A comprehensive evaluation of normalization methods for Illumina high-throughput RNA sequencing data analysis. Brief Bioinform 14(6), 671–683. doi: 10.1093/bib/bbs046.

Escorcia-Rodriguez, J.M., Tauch, A., and Freyre-Gonzalez, J.A. (2020). Abasy Atlas v2.2: The most comprehensive and up-to-date inventory of meta-curated, historical, bacterial regulatory networks, their completeness and system-level characterization. Comput Struct Biotechnol J 18, 1228–1237. doi: 10.1016/j.csbj.2020.05.015.

Escorcia-Rodriguez, J.M., Tauch, A., and Freyre-Gonzalez, J.A. (2021). Corynebacterium glutamicum Regulation beyond Transcription: Organizing Principles and Reconstruction of an Extended Regulatory Network Incorporating Regulations Mediated by Small RNA and Protein-Protein Interactions. Microorganisms 9(7). doi: 10.3390/microorganisms9071395.

Evans, C., Hardin, J., and Stoebel, D.M. (2018). Selecting between-sample RNA-Seq normalization methods from the perspective of their assumptions. Brief Bioinform 19(5), 776–792. doi: 10.1093/bib/bbx008.

Faith, J.J., Hayete, B., Thaden, J.T., Mogno, I., Wierzbowski, J., Cottarel, G., et al. (2007). Large-scale mapping and validation of Escherichia coli transcriptional regulation from a compendium of expression profiles. PLoS Biol 5(1), e8. doi: 10.1371/journal.pbio.0050008.

Freyre-Gonzalez, J.A., Alonso-Pavon, J.A., Trevino-Quintanilla, L.G., and Collado-Vides, J. (2008). Functional architecture of Escherichia coli: new insights provided by a natural decomposition approach. Genome Biol 9(10), R154. doi: 10.1186/gb-2008-9-10-r154.

Freyre-Gonzalez, J.A., Escorcia-Rodriguez, J.M., Gutierrez-Mondragon, L.F., Marti-Vertiz, J., Torres-Franco, C.N., and Zorro-Aranda, A. (2022). System Principles Governing the Organization, Architecture, Dynamics, and Evolution of Gene Regulatory Networks. Front Bioeng Biotechnol 10, 888732. doi: 10.3389/fbioe.2022.888732.

Freyre-Gonzalez, J.A., and Tauch, A. (2017). Functional architecture and global properties of the Corynebacterium glutamicum regulatory network: Novel insights from a dataset with a high genomic coverage. J Biotechnol 257, 199–210. doi: 10.1016/j.jbiotec.2016.10.025.

Freyre-Gonzalez, J.A., Trevino-Quintanilla, L.G., Valtierra-Gutierrez, I.A., Gutierrez-Rios, R.M., and Alonso-Pavon, J.A. (2012). Prokaryotic regulatory systems biology: Common principles governing the functional architectures of Bacillus subtilis and Escherichia coli unveiled by the natural decomposition approach. J Biotechnol 161(3), 278–286. doi: 10.1016/j.jbiotec.2012.03.028.

Giorgi, F.M., Del Fabbro, C., and Licausi, F. (2013). Comparative study of RNA-seq- and microarray-derived coexpression networks in Arabidopsis thaliana. Bioinformatics 29(6), 717–724. doi: 10.1093/bioinformatics/btt053.

Haury, A.C., Mordelet, F., Vera-Licona, P., and Vert, J.P. (2012). TIGRESS: Trustful Inference of Gene REgulation using Stability Selection. BMC Syst Biol 6, 145. doi: 10.1186/1752-0509-6-145.

Heckathorn, D.D., and Cameron, C.J. (2017). Network Sampling: From Snowball and Multiplicity to Respondent-Driven Sampling. Annual Review of Sociology 43(1), 101–119. doi: 10.1146/annurev-soc-060116-053556.

Hecker, M., Lambeck, S., Toepfer, S., van Someren, E., and Guthke, R. (2009). Gene regulatory network inference: data integration in dynamic models-a review. Biosystems 96(1), 86–103. doi: 10.1016/j.biosystems.2008.12.004.

Huynh-Thu, V.A., Irrthum, A., Wehenkel, L., and Geurts, P. (2010). Inferring regulatory networks from expression data using tree-based methods. PLoS One 5(9). doi: 10.1371/journal.pone.0012776.

Iancu, O.D., Kawane, S., Bottomly, D., Searles, R., Hitzemann, R., and McWeeney, S. (2012). Utilizing RNA-Seq data for de novo coexpression network inference. Bioinformatics 28(12), 1592–1597. doi: 10.1093/bioinformatics/bts245.

Imbert, A., Valsesia, A., Le Gall, C., Armenise, C., Lefebvre, G., Gourraud, P.A., et al. (2018). Multiple hot-deck imputation for network inference from RNA sequencing data. Bioinformatics 34(10), 1726–1732. doi: 10.1093/bioinformatics/btx819.

Johnson, W.E., Li, C., and Rabinovic, A. (2007). Adjusting batch effects in microarray expression data using empirical Bayes methods. Biostatistics 8(1), 118–127. doi: 10.1093/biostatistics/kxj037.

Kim, M., Rai, N., Zorraquino, V., and Tagkopoulos, I. (2016). Multi-omics integration accurately predicts cellular state in unexplored conditions for Escherichia coli. Nat Commun 7, 13090. doi: 10.1038/ncomms13090.

Kuffner, R., Petri, T., Tavakkolkhah, P., Windhager, L., and Zimmer, R. (2012). Inferring gene regulatory networks by ANOVA. Bioinformatics 28(10), 1376–1382. doi: 10.1093/bioinformatics/bts143.

Larsen, S.J., Rottger, R., Schmidt, H., and Baumbach, J. (2019). E. coli gene regulatory networks are inconsistent with gene expression data. Nucleic Acids Res 47(1), 85–92. doi: 10.1093/nar/gky1176.

Lim, H.G., Rychel, K., Sastry, A.V., Bentley, G.J., Mueller, J., Schindel, H.S., et al. (2022). Machine-learning from Pseudomonas putida KT2440 transcriptomes reveals its transcriptional regulatory network. Metab Eng 72, 297–310. doi: 10.1016/j.ymben.2022.04.004.

Lo, K., Raftery, A.E., Dombek, K.M., Zhu, J., Schadt, E.E., Bumgarner, R.E., et al. (2012). Integrating external biological knowledge in the construction of regulatory networks from time-series expression data. BMC Syst Biol 6, 101. doi: 10.1186/1752-0509-6-101.

Love, M.I., Huber, W., and Anders, S. (2014). Moderated estimation of fold change and dispersion for RNA-seq data with DESeq2. Genome Biol 15(12), 550. doi: 10.1186/s13059-014-0550-8.

Lowe, R., Shirley, N., Bleackley, M., Dolan, S., and Shafee, T. (2017). Transcriptomics technologies. PLoS Comput Biol 13(5), e1005457. doi: 10.1371/journal.pcbi.1005457.

Marbach, D., Costello, J.C., Kuffner, R., Vega, N.M., Prill, R.J., Camacho, D.M., et al. (2012). Wisdom of crowds for robust gene network inference. Nat Methods 9(8), 796–804. doi: 10.1038/nmeth.2016.

Marbach, D., Prill, R.J., Schaffter, T., Mattiussi, C., Floreano, D., and Stolovitzky, G. (2010). Revealing strengths and weaknesses of methods for gene network inference. Proc Natl Acad Sci U S A 107(14), 6286–6291. doi: 10.1073/pnas.0913357107.

Margolin, A.A., Nemenman, I., Basso, K., Wiggins, C., Stolovitzky, G., Dalla Favera, R., et al. (2006). ARACNE: an algorithm for the reconstruction of gene regulatory networks in a mammalian cellular context. BMC Bioinformatics 7 Suppl 1, S7. doi: 10.1186/1471-2105-7-S1-S7.

Maza, E., Frasse, P., Senin, P., Bouzayen, M., and Zouine, M. (2013). Comparison of normalization methods for differential gene expression analysis in RNA-Seq experiments: A matter of relative size of studied transcriptomes. Commun Integr Biol 6(6), e25849. doi: 10.4161/cib.25849.

Meyer, P.E., Kontos, K., Lafitte, F., and Bontempi, G. (2007). Information-theoretic inference of large transcriptional regulatory networks. EURASIP J Bioinform Syst Biol, 79879. doi: 10.1155/2007/79879.

Michoel, T., De Smet, R., Joshi, A., Van de Peer, Y., and Marchal, K. (2009). Comparative analysis of module-based versus direct methods for reverse-engineering transcriptional regulatory networks. BMC Syst Biol 3, 49. doi: 10.1186/1752-0509-3-49.

Parise, D., Parise, M.T.D., Kataka, E., Kato, R.B., List, M., Tauch, A., et al. (2021). On the Consistency between Gene Expression and the Gene Regulatory Network of Corynebacterium glutamicum. Netw Syst Med 4(1), 51–59. doi: 10.1089/nsm.2020.0014.

Pratapa, A., Jalihal, A.P., Law, J.N., Bharadwaj, A., and Murali, T.M. (2020). Benchmarking algorithms for gene regulatory network inference from single-cell transcriptomic data. Nat Methods 17(2), 147–154. doi: 10.1038/s41592-019-0690-6.

Proost, S., Krawczyk, A., and Mutwil, M. (2017). LSTrAP: efficiently combining RNA sequencing data into co-expression networks. BMC Bioinformatics 18(1), 444. doi: 10.1186/s12859-017-1861-z.

Saito, T., and Rehmsmeier, M. (2015). The precision-recall plot is more informative than the ROC plot when evaluating binary classifiers on imbalanced datasets. PLoS One 10(3), e0118432. doi: 10.1371/journal.pone.0118432.

Salleh, S.M., Mazzoni, G., Lovendahl, P., and Kadarmideen, H.N. (2018). Gene co-expression networks from RNA sequencing of dairy cattle identifies genes and pathways affecting feed efficiency. BMC Bioinformatics 19(1), 513. doi: 10.1186/s12859-018-2553-z.

Salzberg, S.L. (2019). Next-generation genome annotation: we still struggle to get it right. Genome Biol 20(1), 92. doi: 10.1186/s13059-019-1715-2.

Sastry, A.V., Gao, Y., Szubin, R., Hefner, Y., Xu, S., Kim, D., et al. (2019). The Escherichia coli transcriptome mostly consists of independently regulated modules. Nat Commun 10(1), 5536. doi: 10.1038/s41467-019-13483-w.

Schaffter, T., Marbach, D., and Floreano, D. (2011). GeneNetWeaver: in silico benchmark generation and performance profiling of network inference methods. Bioinformatics 27(16), 2263–2270. doi: 10.1093/bioinformatics/btr373.

Schieber, T.A., Carpi, L., Diaz-Guilera, A., Pardalos, P.M., Masoller, C., and Ravetti, M.G. (2017). Quantification of network structural dissimilarities. Nat Commun 8, 13928. doi: 10.1038/ncomms13928.

Secilmis, D., Hillerton, T., Tjarnberg, A., Nelander, S., Nordling, T.E.M., and Sonnhammer, E.L.L. (2022). Knowledge of the perturbation design is essential for accurate gene regulatory network inference. Sci Rep 12(1), 16531. doi: 10.1038/s41598-022-19005-x.

Sirbu, A., Ruskin, H.J., and Crane, M. (2010). Cross-platform microarray data normalisation for regulatory network inference. PLoS One 5(11), e13822. doi: 10.1371/journal.pone.0013822.

Smid, M., Coebergh van den Braak, R.R.J., van de Werken, H.J.G., van Riet, J., van Galen, A., de Weerd, V., et al. (2018). Gene length corrected trimmed mean of M-values (GeTMM) processing of RNA-seq data performs similarly in intersample analyses while improving intrasample comparisons. BMC Bioinformatics 19(1), 236. doi: 10.1186/s12859-018-2246-7.

Soneson, C., and Delorenzi, M. (2013). A comparison of methods for differential expression analysis of RNA-seq data. BMC Bioinformatics 14, 91. doi: 10.1186/1471-2105-14-91.

Stolovitzky, G., Prill, R.J., and Califano, A. (2009). Lessons from the DREAM2 Challenges. Ann N Y Acad Sci 1158, 159–195. doi: 10.1111/j.1749-6632.2009.04497.x.

Swarbreck, D., Wilks, C., Lamesch, P., Berardini, T.Z., Garcia-Hernandez, M., Foerster, H., et al. (2008). The Arabidopsis Information Resource (TAIR): gene structure and function annotation. Nucleic Acids Res 36(Database issue), D1009–1014. doi: 10.1093/nar/gkm965.

Taboada-Castro, H., Gil, J., Gomez-Caudillo, L., Escorcia-Rodriguez, J.M., Freyre-Gonzalez, J.A., and Encarnacion-Guevara, S. (2022). Rhizobium etli CFN42 proteomes showed isoenzymes in free-living and symbiosis with a different transcriptional regulation inferred from a transcriptional regulatory network. Front Microbiol 13, 947678. doi: 10.3389/fmicb.2022.947678.

Van den Bulcke, T., Van Leemput, K., Naudts, B., van Remortel, P., Ma, H., Verschoren, A., et al. (2006). SynTReN: a generator of synthetic gene expression data for design and analysis of structure learning algorithms. BMC Bioinformatics 7, 43. doi: 10.1186/1471-2105-7-43.

Young, W.C., Raftery, A.E., and Yeung, K.Y. (2014). Fast Bayesian inference for gene regulatory networks using ScanBMA. BMC Syst Biol 8, 47. doi: 10.1186/1752-0509-8-47.

Zhang, B., and Horvath, S. (2005). A general framework for weighted gene co-expression network analysis. Stat Appl Genet Mol Biol 4, Article17. doi: 10.2202/1544-6115.1128.

Zhang, J., Nie, Q., Si, C., Wang, C., Chen, Y., Sun, W., et al. (2019). Weighted Gene Co-expression Network Analysis for RNA-Sequencing Data of the Varicose Veins Transcriptome. Front Physiol 10, 278. doi: 10.3389/fphys.2019.00278.

Zorro-Aranda, A., Escorcia-Rodriguez, J.M., Gonzalez-Kise, J.K., and Freyre-Gonzalez, J.A. (2022). Curation, inference, and assessment of a globally reconstructed gene regulatory network for Streptomyces coelicolor. Sci Rep 12(1), 2840. doi: 10.1038/s41598-022-06658-x.

