## Supplementary File 2 for "Improving gene regulatory network inference and assessment: The importance of using network structure"

Supplementary material

### ∑ Background

#### Regression

Most of the methods based on regression aim to directly predict the behavior of a target gene given the expression pattern of a transcription factor (TF). There are two main regression methodologies for these inference methods: Least Angle Regression (LARS) and Least Absolute Shrinkage and Selection Operator (LASSO). In general, both algorithms reduce the number of TFs necessary to explain the behavior of one gene.

##### Trustful Inference of Gene Regulation using Stability Selection (TIGRESS) (CAUS)

TIGRESS is based on LARS and “stability selection”; each iteration runs the LARS algorithm and then re-samples the genes and TFs. TIGRESS predicts all interactions between each TF and all the other genes and weighs them based on the frequency of the chosen TF as a regulator of the gene among all iterations. This inference method takes as input a list of putative TFs and a gene expression matrix and, the output is a list of weighted interactions of the directed network. Although TIGRESS assumes there will be more genes than conditions, the number of TFs used cannot be greater than the number of conditions. Also, the expression data must have been pre-processed for quality control and missing values must be imputed (Haury et al.). TIGRESS is available online, alongside proper documentation and examples. TIGRESS was ranked among the top inference method in the DREAM5 gene network inference challenge and it was awarded as the best linear regression-based method in the assessment (Marbach et al.). TIGRESS has also been assessed with Drosophila melanogaster gene expression data (Kang et al.). Like other regression-based methods, TIGRESS ignores the influences from unknown or unobserved regulators, which degrades its performance.

##### Statmodel (CAUS)

Statmodel is a method for Statistical Modeling and Analysis of Experiments without ANOVA. It aims to achieve a more dependable real-world experimental analysis avoiding the many requirements of ANOVA, which if they are not fulfilled, could cause misled conclusions. In this method, the test of the hypothesis required by the statistical analysis is performed using optimal parameter identification. After some mathematical treatment of the data, it performs a hypothesis test applying a $c^{2}$ test in Matlab, to test the influence of a variable For the GRN inference, a TF, is significantly different from zero on the response variable, in the case of GCN inference, a TG. The score of the interaction is proposed as the $-\log\left( p-value \right)$ of the hypothesis test (Hernandez). The algorithm takes as input a matrix of expression data and a list of TFs, and the output is a directed network into the list format of TF-TG interactions and their correlation coefficients.

#### Ensemble Learning

Ensemble methods perform feature selection combining multiple machine learning algorithms, such as filter methods, wrapper methods, and embedded methods; to decrease variance, bias and improve predictions, often obtaining better performance than when applying a single machine learning algorithm (Ruyssinck et al.).

##### GEne Network Inference with Ensemble of trees (GENIE3) (CAUS)

This method is based on two main frameworks: regression and feature selection. First, it breaks down the problem into n regression problems, with n being equal to the number of genes. In each regression problem, the expression of a target gene is predicted by considering all other genes as candidate regulators. Then, using feature selection, it is determined which of the candidate regulators are the most relevant for predicting the target gene expression profile. GENIE3 ranks the interactions obtained using a tree-based regression model doing a global ranking of all the network interactions (Huynh-Thu et al.). GENIE3 has been implemented in MATLAB, Python, and R programming languages, each one with different running times, highlighting the R wrapper of a C code for being the fastest. The documentation of the software is very concrete and contains tutorials with its own data, which allows the user to easily follow the steps in the process. Also, on the same website, it is possible to download other GRN inference algorithms developed by the group. In our work, we used its MATLAB implementation. GENIE3 has been a subject of comparison for many recent publications of GRN inference algorithms. Its fame is due to its recognition by the DREAM5 consortium as the best predictor in two of their competitions; quickly becoming a state-of-the-art algorithm for GRN inference.

#### Meta

Meta methods are strategies that combine diverse inference approximation, like regression, correlation, etc. for gene network inference.

##### Inferelator (CAUS)

Inferelator is an algorithm that uses standard regression and model shrinkage techniques to select parsimonious, predictive models for the expression of a gene, or cluster of genes, as a function of the levels of the transcription factors, environmental influences, and interactions between these factors. As input, it requires an expression matrix that can be time-series or steady-state, where the rows are genes and the columns are the conditions. The method’s performance can be improved by reducing the dimensionality of the data before the inferring process. This can be done by specifying a list of regulators and performing a biclustering method to the expression matrix, for which the authors recommend using cMonkey. On their webpage, you can find several versions, as well as minimal instructions for the installation and execution of the algorithm. However, if there are complications along the process, the authors have a google group with the purpose to help the users. In our work we used Inferelator_2015.03.03. The required input consists of a matrix where columns are experiments and rows are genes, as well as a list of TFs. The output consists of a list of directed regulatory interactions (Bonneau et al.).

#### Bayesian

A Bayesian network is a directed acyclic graph, where each link represents the probability distribution for the gene A regulating a gene B. Defining father-son relationships between genes, where a descendant cannot regulate its own ancestor. The Bayesian network inference methods are characterized by the optional incorporation of prior biological knowledge, which leads to a better output network. The major limitations for the Bayesian inference methods are the computational cost and the requirement of large datasets for generating the initial step of the algorithm (Chai et al.; Linde et al.). However, these methods can incorporate interactions previously known and biological data from diverse sources (Linde et al.). Also, during the DREAM4 competition, it was highlighted that Bayesian methods have an overall deficient performance predicting fan-in and fan-out motifs; while their predictions of cascade motifs are the most accurate (Marbach et al.).

##### iterativeBMA (CAUS)

To apply BMA to several potential regulators, the authors have developed iterative BMA (iBMA). First, a linear model is fitted such that the expression of candidate regulators from a previous time point must be able to predict the expression level of a target gene in the current time point; variables are ranked in order of their determination coefficients from fitting single variable models. Then, BMA is applied to the top 30 genes. Candidate regulators with the lowest assigned posterior probabilities are removed. Finally, an equal number of candidates from the original ranked list are added to apply BMA to 30 genes once again. This process is continued until the top 100 genes are considered (Lo et al.).

##### ScanBMA (CAUS)

As part of the BMA family, this method builds the network from a variable selection approach, in which the regulators of each gene are inferred. Using a greedy algorithm and Occam’s window principle, ScanBMA can reduce the space of models of the BMA methodology that would be used to compute the posterior probabilities by excluding the unlikely models. Additionally, Zellner’s g-prior is used to get the prior distributions of the model parameters and the excess of variation is reduced from the expression data by removing the effects that the genes could have on themselves hence improving the accuracy of the inferred edges. The assessment of this method by the authors consisted of the inference of two datasets, one from yeast data and another simulated gene expression data, which resulted in better performance or comparably to other GRN inference methods depending on the prior information availability (Young et al.).

#### Correlation

The principle of correlation inference methods states that if two genes participate in a regulatory interaction, they have correlated expression profiles. It is important to remark that gene expression correlation is a necessary, but not a sufficient condition to infer a network interaction.

##### Large-Scale Transcriptome Analysis Pipeline (LSTrAP) (COEX)

LSTrAP combines different tools to process data from RNA-seq expression profiles. The main goal of LSTrAP is to infer a gene network efficiently by generating a pipeline. This pipeline identifies the low-quality data, administrates the computational resources, and generates the correlation network between gene profiles. The documentation is complete and detailed guiding the user step by step and providing tools for the pre- and post-processing of the data. LSTrAP requires as input the gene expression as fastq files and the genome of the organism. The output is not just the coexpression network, but also the normalized expression profile, co-expression clusters, and optionally, the protein domains and the gene families by detection of orthologous genes. Additionally, it provides quality control to identify the "potentially problematic" samples. The authors have evaluated their method using RNA-seq data from *Arabidopsis thaliana* and *Sorghum bicolor* to assess how well the pipeline can group genes, in this case, photosynthesis genes (Proost et al.). Since most of the workflow is for processing RNA-seq data, only the Python script that computes the correlations between gene expression profiles was used in this study (pcc.py).

##### Weighted Gene Co-Expression Network Analysis (WGCNA) (COEX)

WGCNA calculates a weighted adjacency matrix based on the similarity between the gene expression profiles. This matrix is then optimized by an adjacency function that maximizes the similitude of the final network to a scale-free topology. As a downstream analysis, the network is partitioned into modules of genes with similar expression profiles, which are also able to correlate additional biological data from the sample to the modules. To infer a complete GRN, WGCNA takes as input the gene expression data matrix and a lambda parameter for the adjacency function. The method assumes that the gene expression data has been suitably quantified and normalized. WGCNA is implemented as an R package with extensive documentation coupled with tutorials available at the author’s website (Zhang and Horvath).

#### Mutual Information

The mutual information (MI) measures the dependence between two random variables, it relates the entropy reduction from a variable given the state of another. In the context of GRN inference, we consider the gene expression values as variable states. The MI between a pair of genes is symmetric, therefore the output is an undirected network (Linde et al., 2015). It is important to notice that given the MI definition, each experiment must be independent of the others which are great for most data sets. However, for time series data this assumption implies that each time point must be separated by enough time to assume its independence (Bansal et al.). The evaluation performed by DREAM4 on the MI predictions highlights their performance detecting non-linear interactions, such as feed-forward loops, whereas the linear interactions are harder to detect (Marbach et al.).

##### Algorithm for the Reconstruction of Accurate Cellular Network (ARACNE) (COEX)

This inference method is performed as a two-step process. First, it infers associations between any two edges in each triplet structure by MI, obtaining a lot of false positive interactions, especially on those probes sharing a similar expression intensity, owes to indirect interaction. Later, data process inequality is used to filter out those indirect interactions. The algorithm is designed to manage the complexity of regulatory networks in mammalian cells, although the authors mention its capability to infer regulatory networks on a wider range of organisms (Margolin et al.). This method has also been implemented as a function in the R package minet (this implementation was used in this study). Additionally, ARACNE is available as a Cytoscape app. The executable files and Java source codes are also available on the website of the laboratory of Andrea Califano. This website also includes the documentation of the method, which is available in both summarized and extended versions. The method takes as input microarray expression profiles to generate an adjacency matrix once the data is processed. It is incapable of processing time-course data. The output network is undirected.

##### C3NET (COEX)

Unlike other methods, C3NET does not aim to infer the complete gene interaction network. Rather, it focuses on predicting what the authors call, the Conservative Causal Core. This core is a marrow of the strongest interactions in the network. To achieve this, C3NET starts by estimating the shared MI between each pair of nodes in the network. The statistical significance of MI values is evaluated using resampling methods to filter out non-significant values. Then, for each node in the network, a single interaction is added between said node and the node with which it shares its maximal pair-wise MI value. The C3NET algorithm is available as an R package. The main input for the inference function is the gene expression data set, where the rows are genes and the columns are different samples. Users may modify several default parameters of the inference function, such as the statistical significance threshold, and the number of iterations performed to get the sampling distribution. The function output is network into a symmetric MI matrix, where non-zero values link two genes. The documentation is clear, concise, and the function parameters are explained in sufficient detail for their use. In this work, we used the c3net R package version 1.1.1. When compared to other inference methods based on estimations of MI (ARACNE, MRNE, RN, and CLR). Because it aims to infer a small number of robust interactions, this method infers fewer interactions than others, having a higher number of true positives (Altay and Emmert-Streib).

##### Context Likelihood of Relatedness (CLR) (COEX)

This method seeks to reduce the true positive-false positive tradeoff that is normally found in methods based on MI. To achieve this, CLR calculates the statistical likelihood of the MI value of a candidate regulator-gene pair, given their network context. The MI value of a regulator-gene pair is compared against the distribution of MI values for all the possible interactions that involve the regulator or the gene. Therefore, the most probable interactions have MI values significantly above the background distribution of MI scores. Consequently, the algorithm removes several false interactions by eliminating cases where a gene weakly (low MI) interacts with several other genes. This can arise due to an indirect interaction, inadequate sampling, or failure in microarray normalization. The CLR algorithm is also available as a function in the minet R package. Just as MRNET, the CLR function receives an MI matrix as input and returns the co-expression network as a weighted adjacency matrix output. CLR has been previously evaluated on E. coli microarray experiments that have been normalized with RMA. Some inferred interactions have been validated experimentally, as well as with binding motif analysis. CLR is used as a comparison for new methods. Some inference methods, even incorporate the CLR algorithm as part of their pipeline (Faith et al.).

##### Minimum Redundancy Networks (MRNET) (COEX)

MRNET is a method that uses the maximum relevance/minimum redundancy (MRMR) algorithm. MRMR is a feature selection algorithm with an information-theoretic approach that measures mutual information to maximize relevance while minimizing redundancy. MRNET selects one gene at a time as the “fixed results” (the independent variable) and the others as the independent features that would predict the result. This procedure is done for each gene. The output of this stage is a weighted adjacency matrix of scores, which then undergoes a pruning phase by selecting a threshold to infer whether there is an interaction or not (Budden and Crampin). MRNET is implemented in the R language, and it is available in the package minet. In our work, we use minet version 3.38.0. The documentation is short, but it is clear enough to use the package. Minet also includes other tools to manipulate the data allowing the user to pre-process it to use the method. The function of the package takes as its main input an MI matrix and outputs a weighted adjacency matrix.

#### Analysis of Variance

Analysis of Variance (ANOVA) is a statistical method used to evaluate the differences between two or more means in a sample. These differences are analyzed through an analysis of variance, discretizing the observed variance into components trackable to diverse sources of variation.

##### ANOVA and Friedman (HYBR)

ANOVA is a method for GRN inference, where the likelihood of an interaction between a transcription factor (TF) and a target gene (TG) is given by a non-linear correlation coefficient, which is derived from an analysis of variance (two-way ANOVA). The correlation coefficient measures association as the fraction of the total variance that is explained by the differential expression across experimental conditions (Kuffner et al.). ANOVA has some requirements for its proper application, one of them is normal distributions. This might not be true in the case of GRN inference. Thus, a variation of the method is proposed, deriving the non-linear correlation coefficient from a Friedman test, which is the non–parametric alternative of ANOVA, since does not make assumptions of normality (Hoffman). The implementation used in this paper, for both of these methods for MATLAB, was not made by Küffner, 2012. The one used in this review can be found at https://github.com/dazorroa/Anova. The algorithm takes as input a matrix of expression data and a list of TFs, and the output is the list of TF - TG correlation coefficients.

### ∑ Results

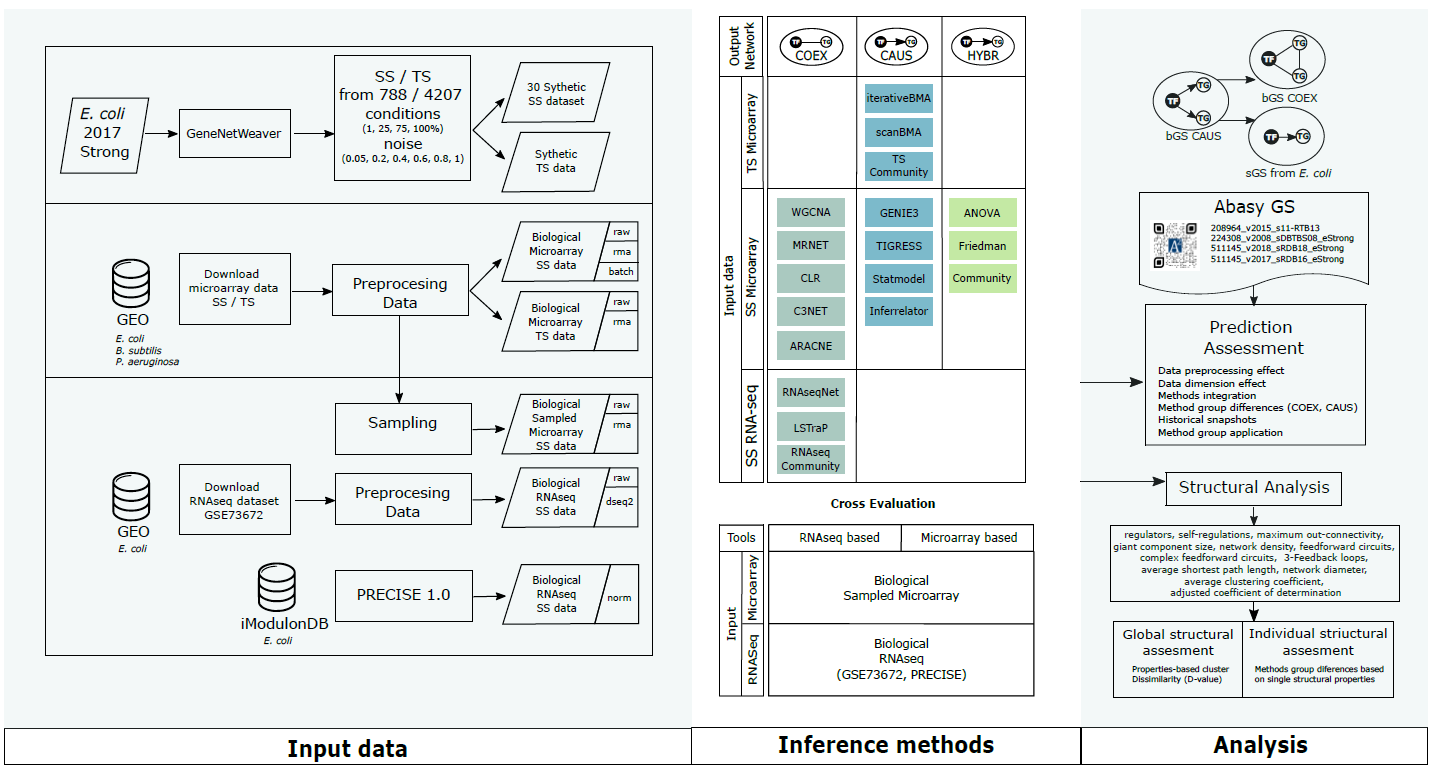

#### Supplementary Figure 1. Detailed pipeline. We assessed the effect related to the properties of genome-wide expression data using: synthetic SS, synthetic TS, biological microarray SS, biological microarray TS, and biological RNA-seq SS. The synthetic data sets were simulated using GeneNetWeber software using the strong *E. coli* GS of Abasy. The biological microarray sets were retrieved for *Escherichia coli* K-12 MG1655, *Bacillus subtilis* 168, and the pathogen *Pseudomonas aeruginosa* PAO1 from GEO. We include a processed RNA-seq dataset PRECISE 1.0. The feasible preprocessing steps were applied according to the dataset (noise %, completeness %, raw, RMA, batch, deseq2, norm). We run all inference tools using the proper input dataset. In addition, we classify inference tools according to the output inferred network (COEX, CAUS, HYBR). Further, we performed a cross-evaluation of tools developed for RNA-seq and for microarray by interchanging input data. Therefore, we assess inference tools' performance in the task of recovering the GSs. Finally, we evaluated inferred networks in terms of structural properties. Static samples (SS), Time-series (TS), non-normalized data (raw), robust multiarray averaging (RMA), batch effect correction (batch), transcript per million normalization (norm), transcription factor (TF), target gene (TG), output network into causal relationship networks (CAUS), co-expression networks (COEX), and hybrid networks (HYBR), gold standard networks (GS), false positive rate (FPR), true positive rate (TPR).

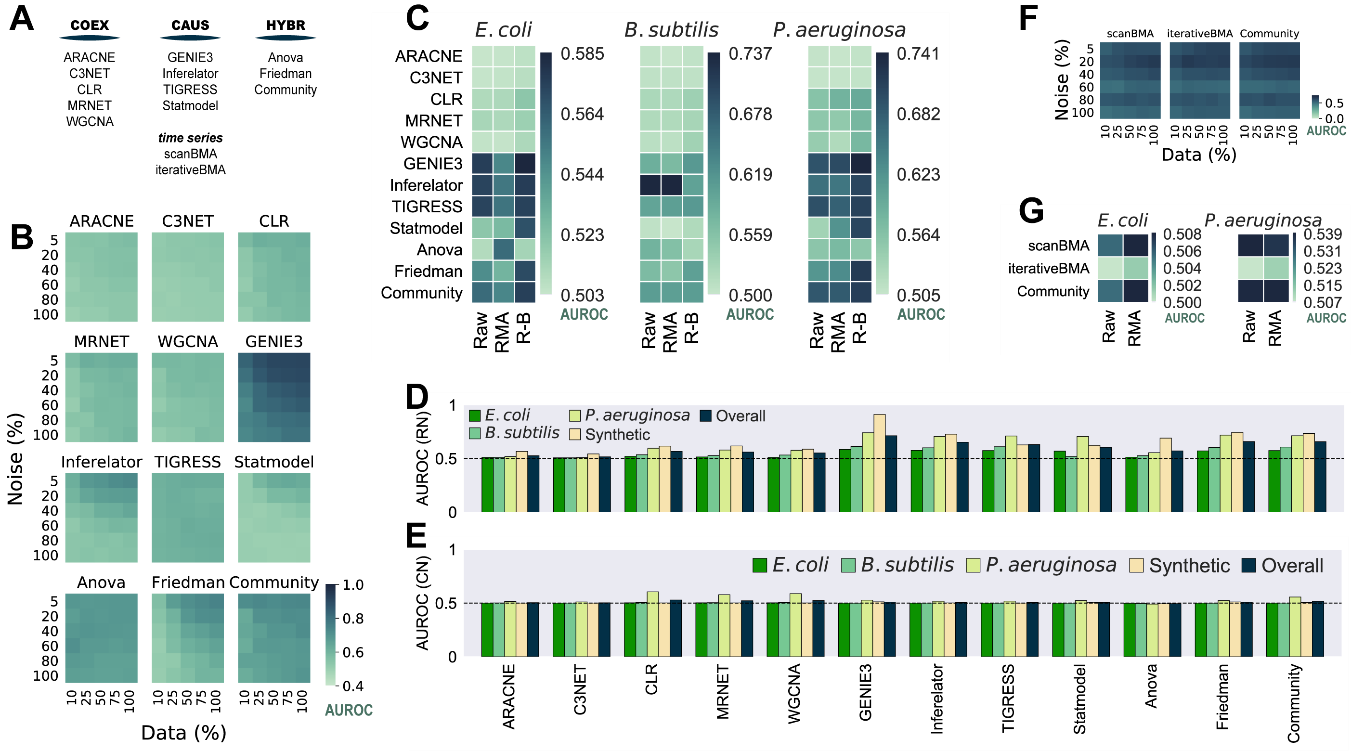

#### Supplementary Figure 2. Assessment using AUROC. Complements Figure 1 in the main text using the area under the ROC (AUROC) curve instead of the area under the precision-recall curve (AUPR).

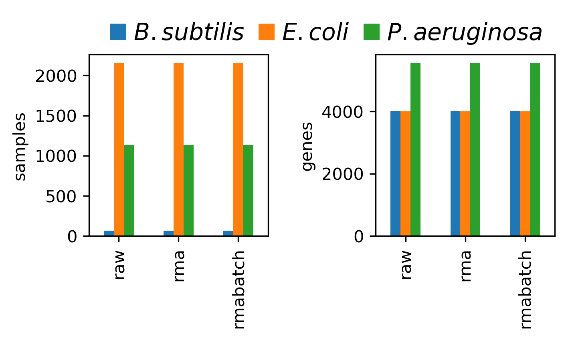

#### Supplementary Figure 3. Data size. *B. subtilis* is the organism with the smaller dataset (64 samples).

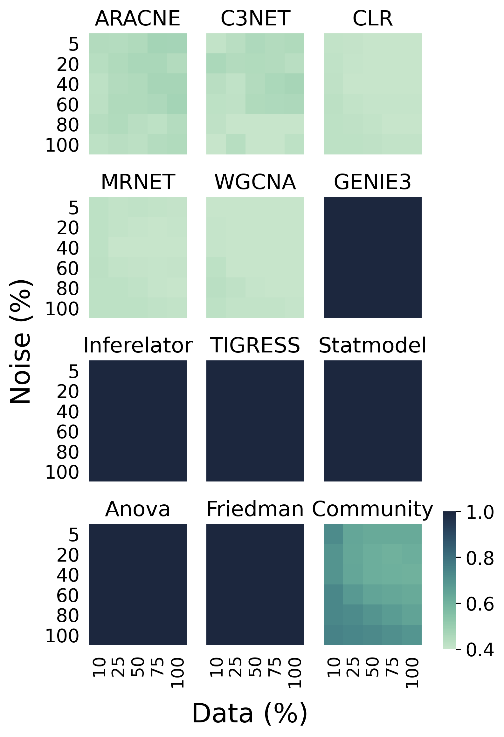

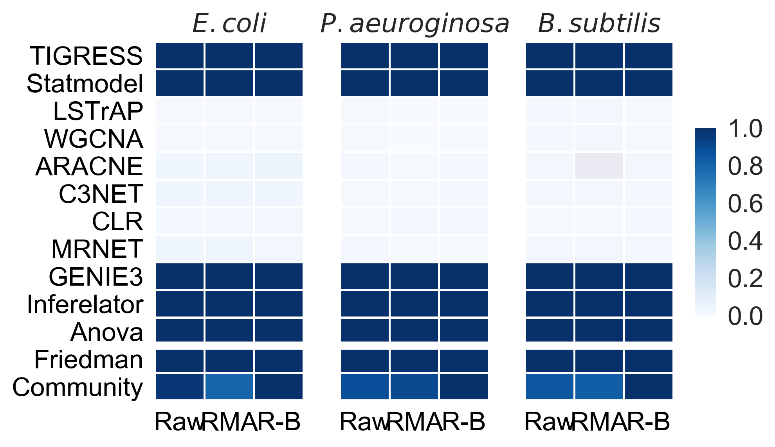

#### Supplementary Figure 4. Fraction of predicted interactions mediated by a known TF in synthetic (left) and biological (right) data. Tools that don’t require the name of the genes coding for TFs are designed to infer co-regulatory networks. For this reason, they have poor performance when assessed against gene regulatory networks.

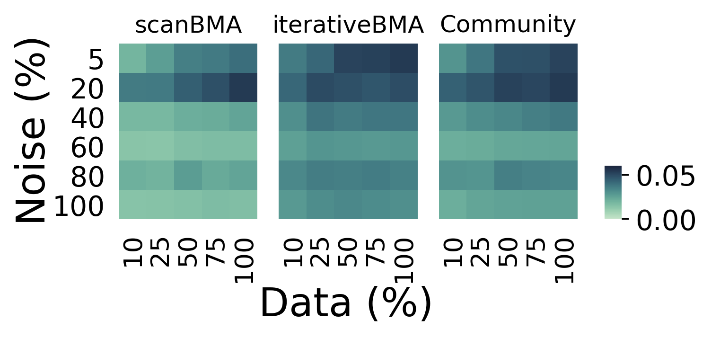

**Supplementary Figure 5.** AUPR with synthetic data for tools working with time series. See Supplementary Figure 2 for AUROC values.

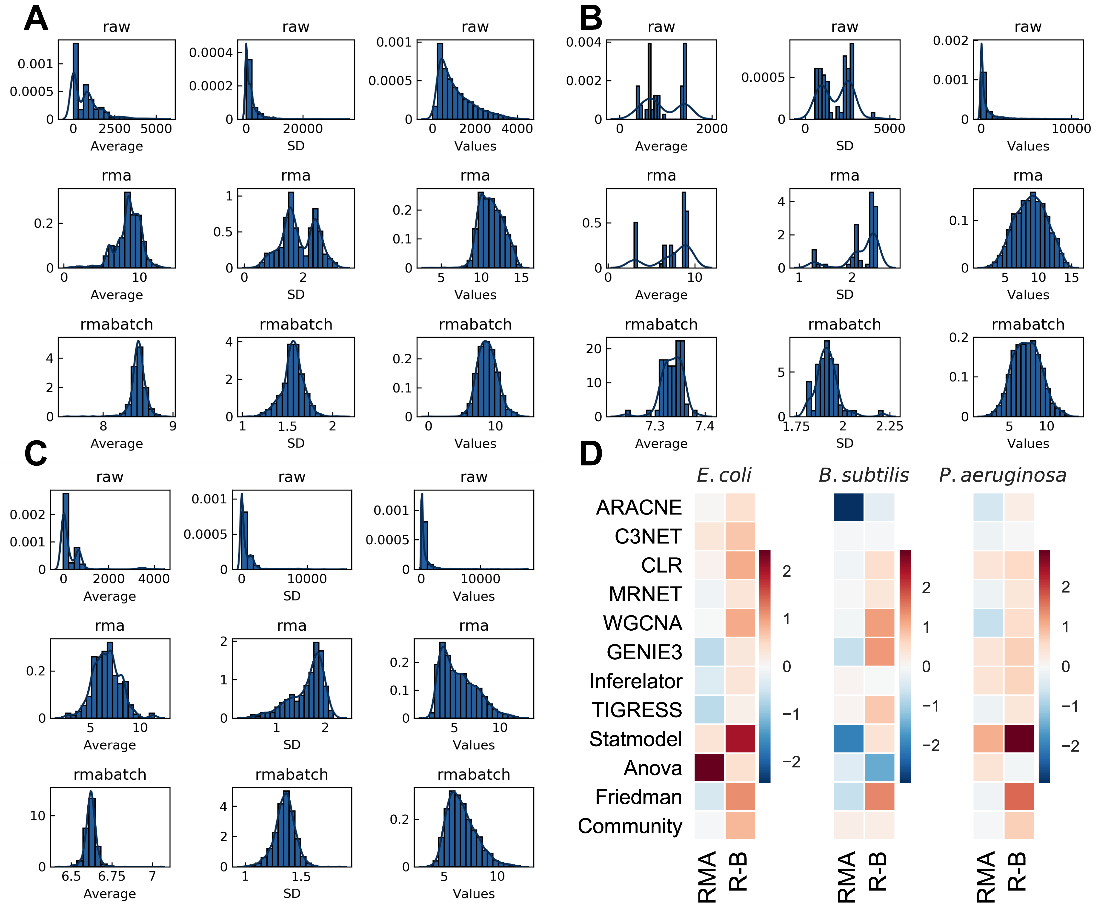

#### Supplementary Figure 6. Effect of the normalization on the datasets for A) *E. coli,* B) *B. subtilis,* and C) *P. aeruginosa.* D) None of the preprocessing techniques seems to have a relevant effect on the AUROC score. RMA normalization and batch correction presents a slight improvement over only RMA normalization.

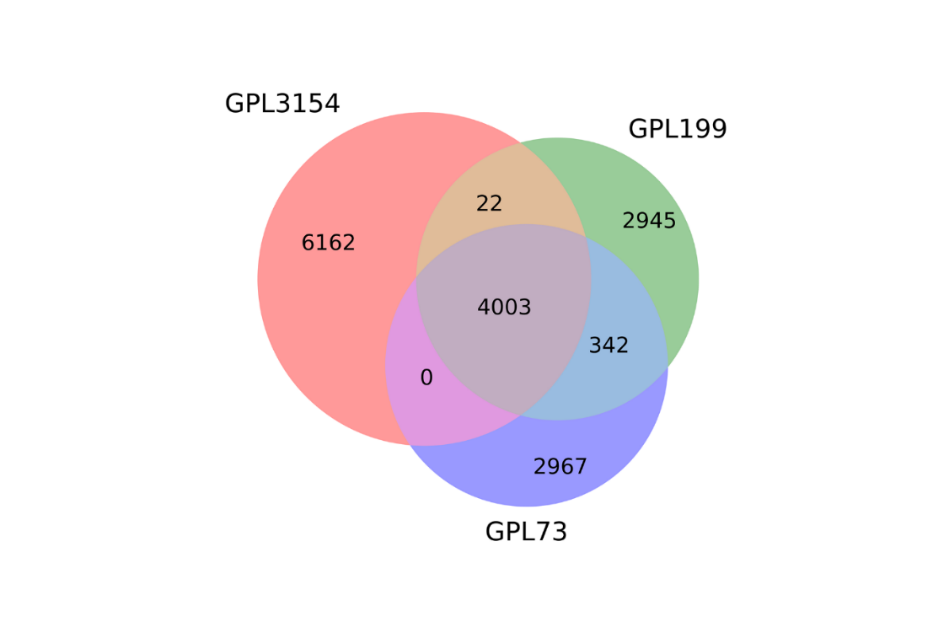

#### Supplementary Figure 7. Common genes of *E. coli* K-12 MG1655 among the platforms. We used the 4003 genes found in the three platforms.

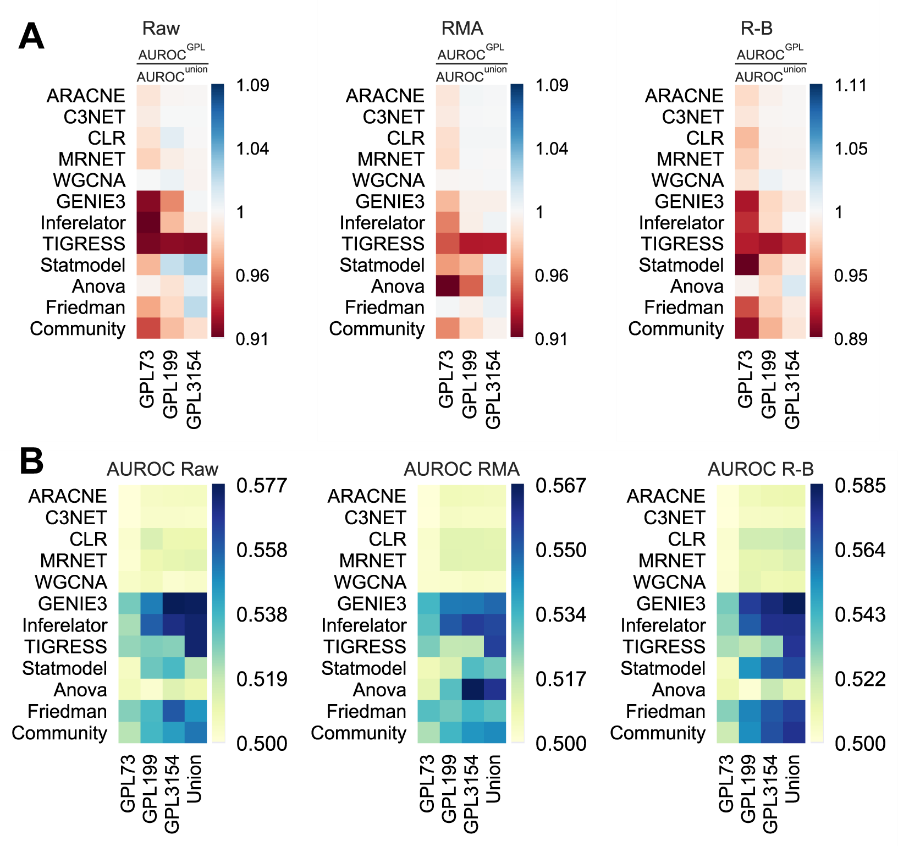

#### Supplementary Figure 8. Methods AUROC score improves with larger datasets as a training set. A) AUROC with reference to the Union. B) Raw values. See figure 3B for size reference.

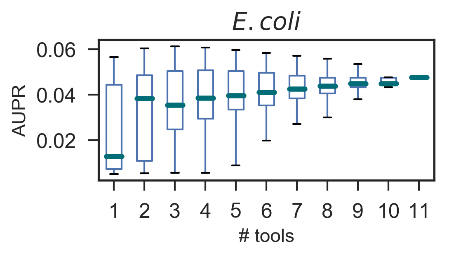

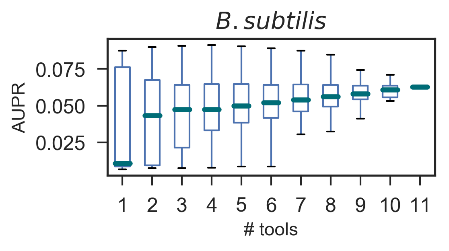

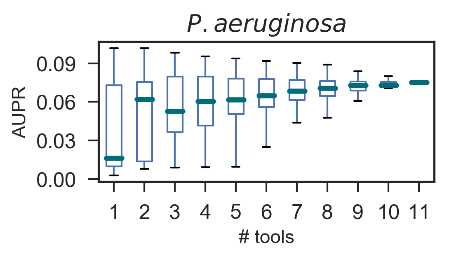

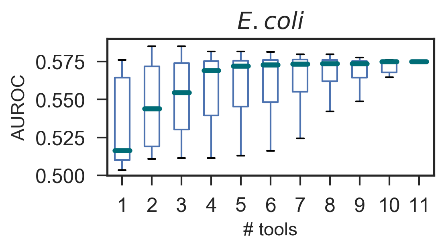

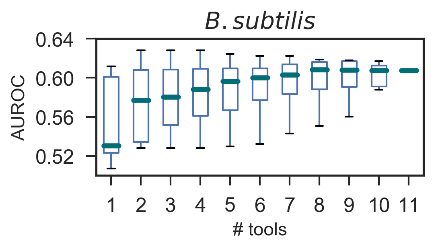

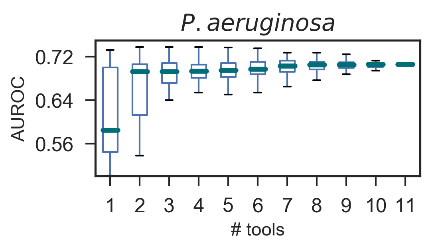

#### Supplementary Figure 9. AUPR and AUROC for the selective community networks.

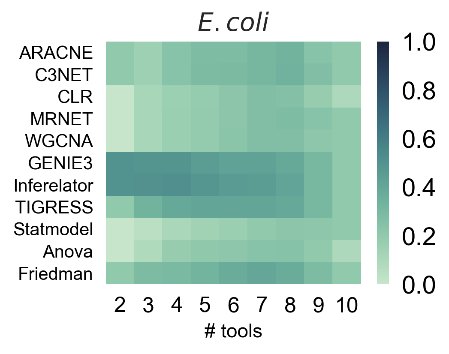

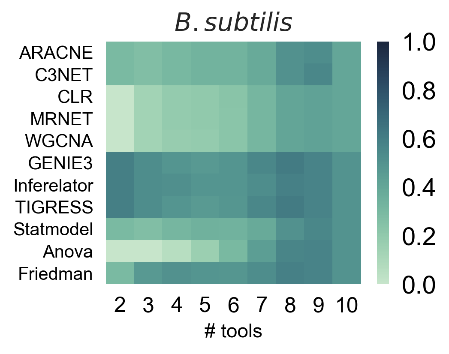

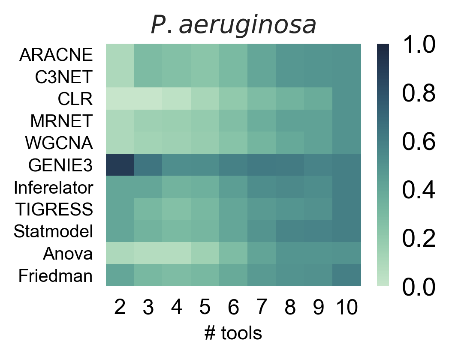

#### Supplementary Figure 10. How the performance of single-tool predictions changes as more predictions are included. The score is the fraction of times a selective community network with the tool corresponding tool (x-axis).

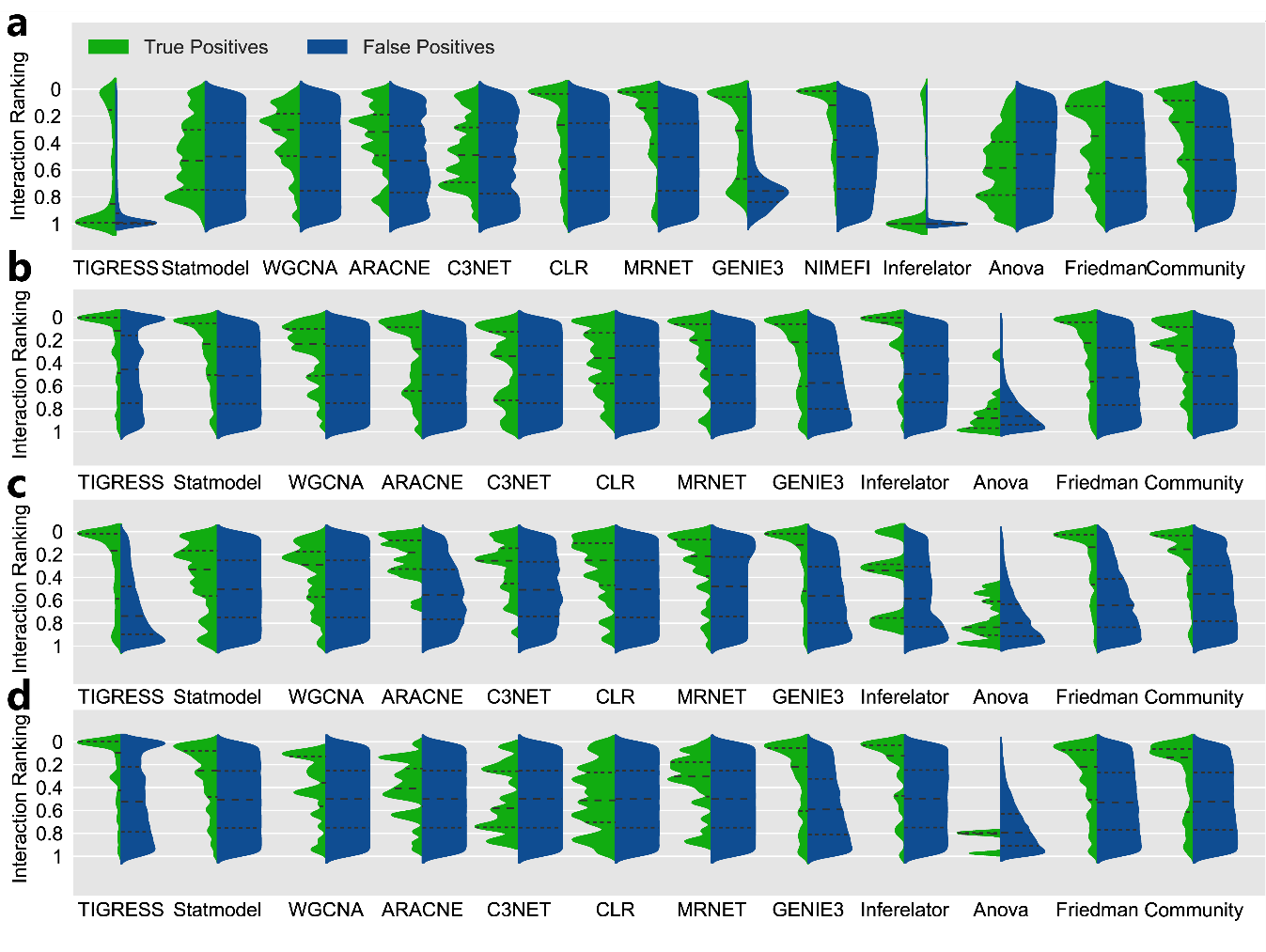

#### Supplementary Figure 11. TP and FP distribution on interaction raking for a) synthetic data, b) *E. coli*, c) *B. subtilis*, and d) *P. aeruginosa* using the first 100000 interactions. The area of the violinplots is not comparable.

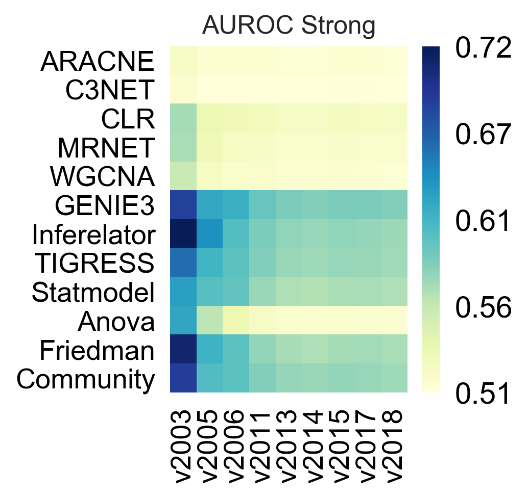

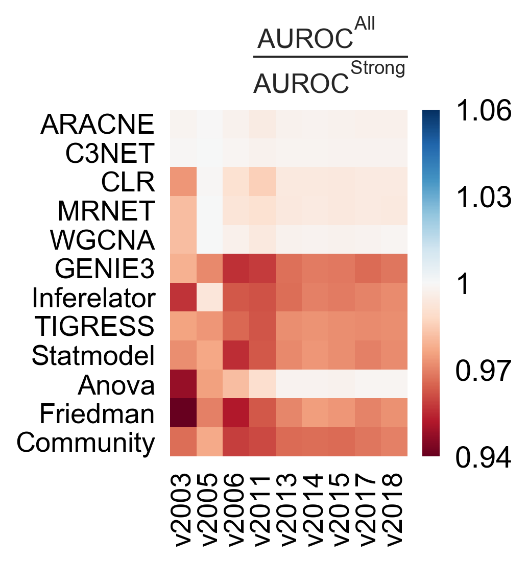

#### Supplementary Figure 12. Assessment with the historical snapshots of *E. coli.*

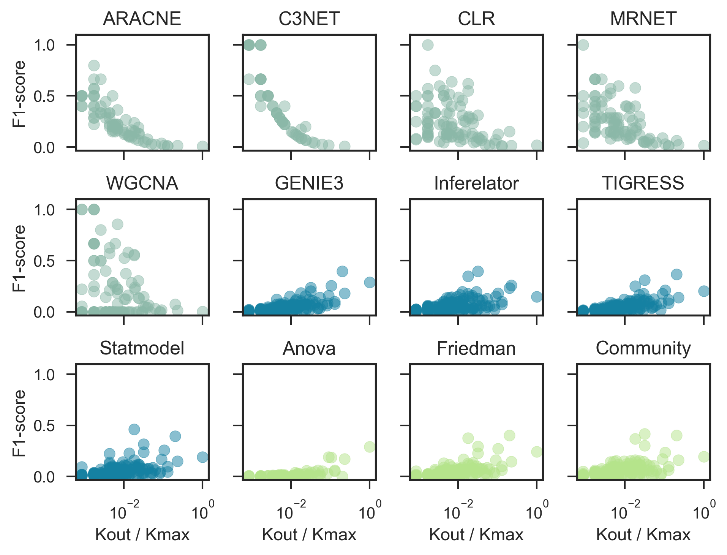

#### Supplementary Figure 13. Regulon level assessment with F1.

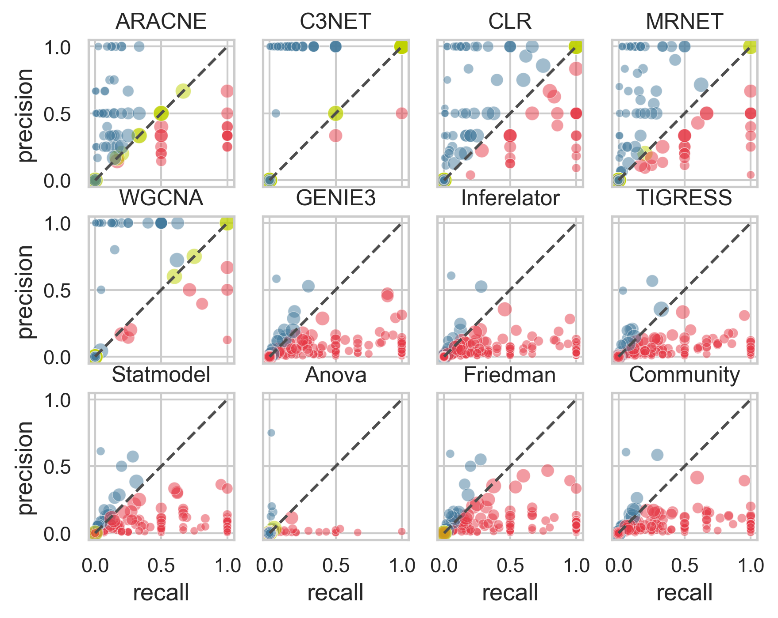

#### Supplementary Figure 14. Sub-(blue) and over(red)-estimation of the number of targets per regulon. There is a dot for each regulon inferred by the tools. Blue dots are inferred regulons sub-estimating the number of targets, while red dots are inferences over-estimating the number of targets for a specific regulon. Green dots placed on the diagonal are for regulons with the same number of targets in the inference and the gold standard. Note that the number might be correct, but the number of targets shaping the regulon is what dictates the dot position. This sub- and over-estimation is also visible analyzing the formulas for precision ( $\frac{\boldsymbol{TP}}{\boldsymbol{TP+FP}}\boldsymbol{)}$ and recall ( $\frac{\boldsymbol{TP}}{\boldsymbol{TP+FN}}\boldsymbol{)}$. Given that $\left| \boldsymbol{Prediction} \right|\boldsymbol{=TP+FP}$ and $\left| \boldsymbol{Gold standard} \right|\boldsymbol{=TP+FN}$, $\boldsymbol{if}\left| \boldsymbol{Prediction} \right|\boldsymbol{=}\left| \boldsymbol{Gold standard} \right|\boldsymbol{\Rightarrow precision=recall}$ (i.e., the diagonal where the number of inferred TGs is the same as the number of TGs in the actual regulon). When there is an over-estimation ($\left| \boldsymbol{Prediction} \right|\boldsymbol{>}\left| \boldsymbol{Gold standard} \right|$) the recall will be higher than the precision. On the other hand, when there is a sub-estimation ($\left| \boldsymbol{Prediction} \right|\boldsymbol{<}\left| \boldsymbol{Gold standard} \right|$) the precision will be higher than the recall.

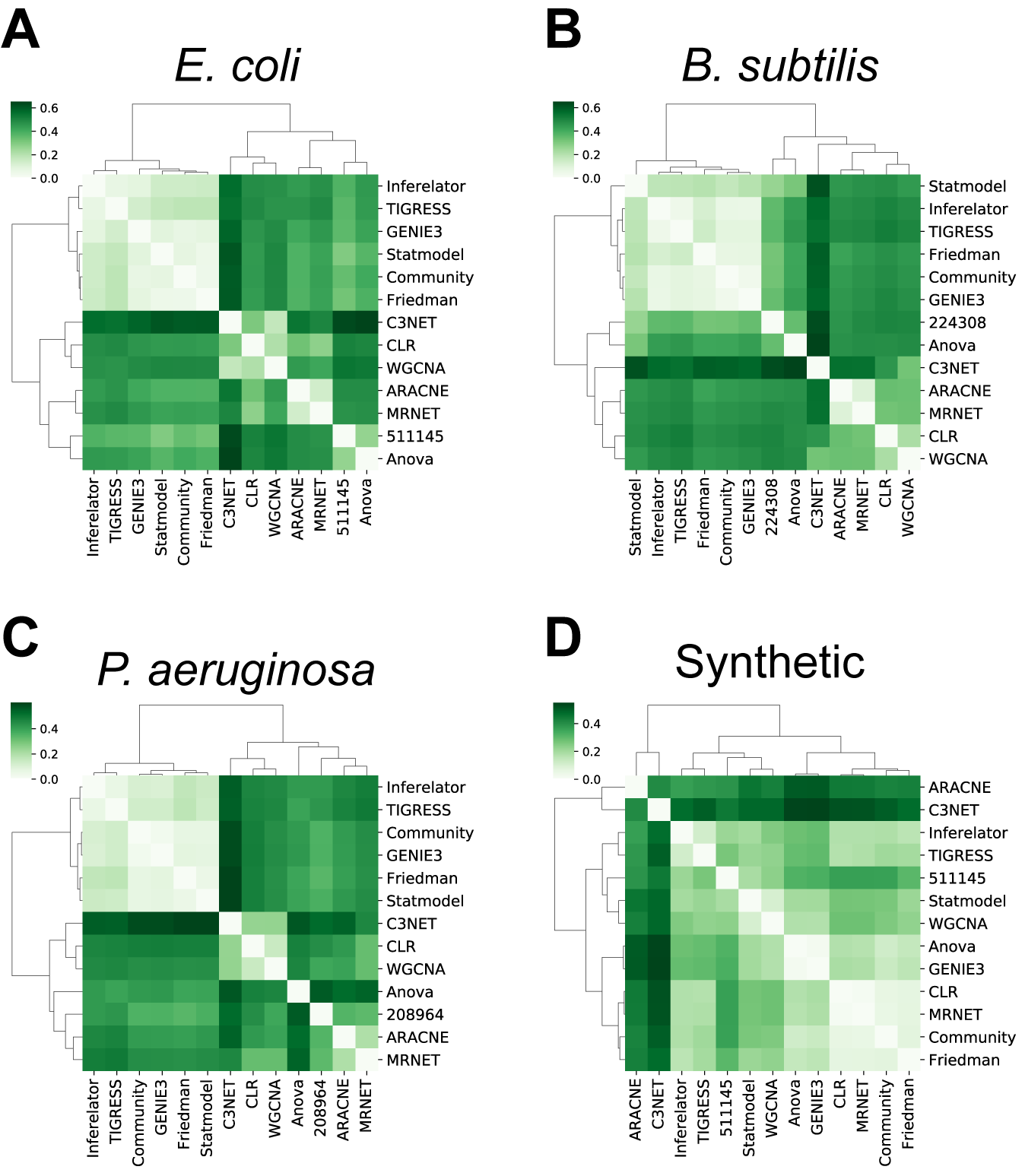

#### Supplementary Figure 15. D-value clustering. The CAUS tools are clustered together along with Community and Friedman with biological data (A-C). Clustering the networks with synthetic data does not identify any of the tool groups according to their type of network. The results agree with the clustering using structural properties.

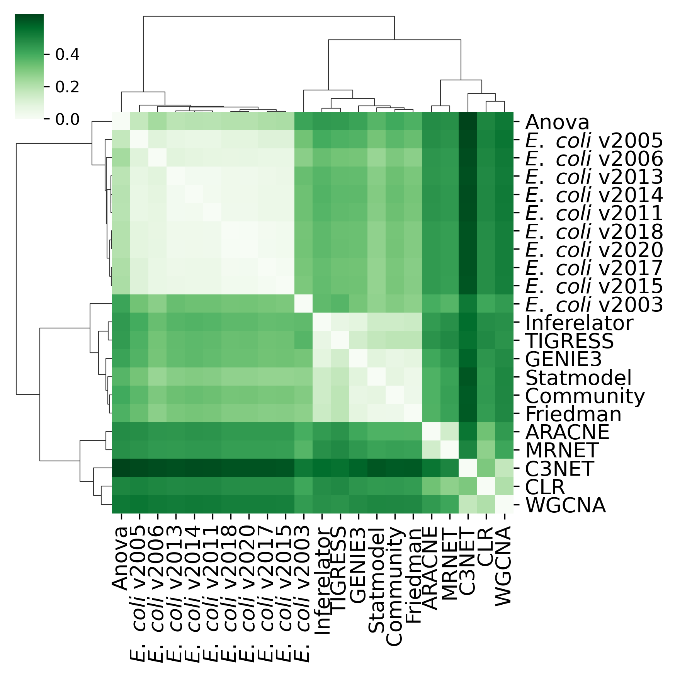

#### Supplementary Figure 16. D-value clustering including the historical snapshots for *E. coli.*

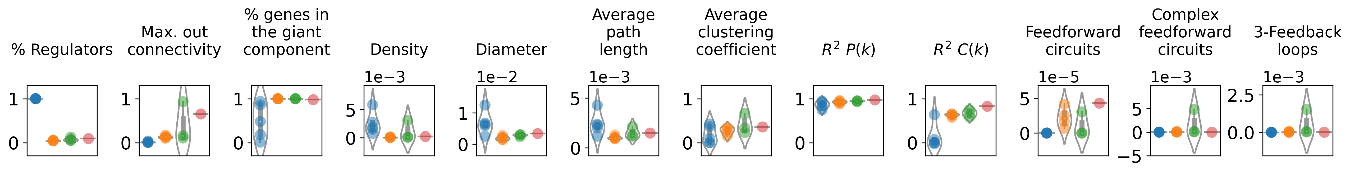

**A**

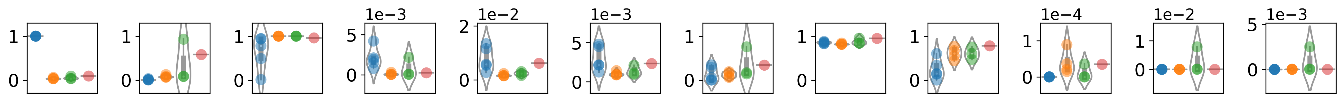
**B**

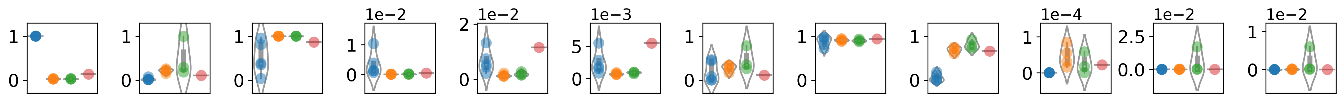
**C**

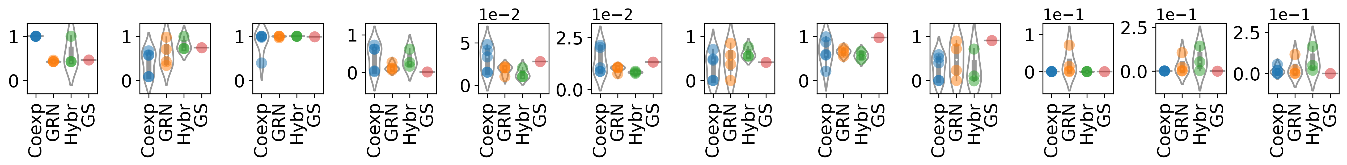
 **D**

#### Supplementary Figure 17. Individual structural properties for A) *E. coli,* B) *B. subtilis,* C) *P. aeruginosa*, and D) synthetic data.

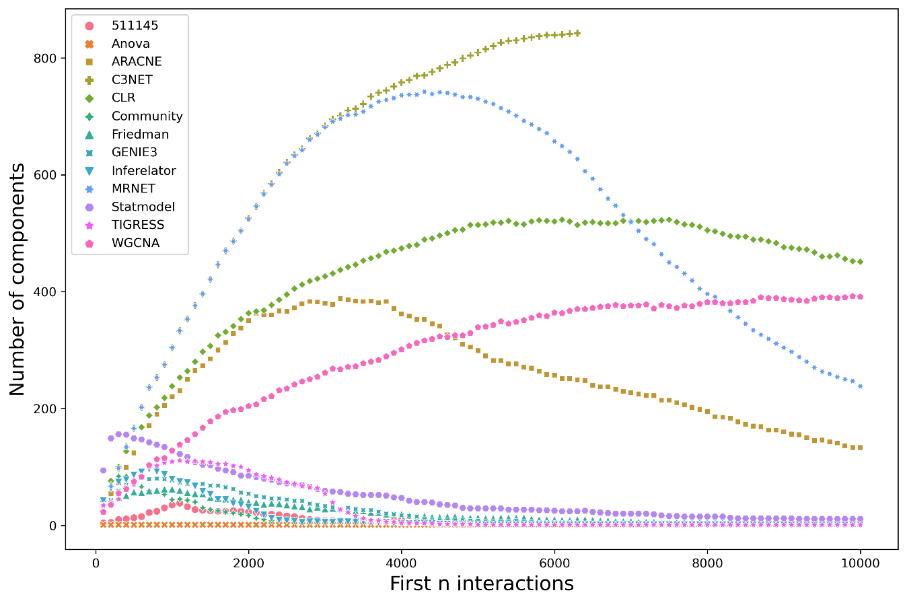

#### Supplementary Figure 18. COEX networks tend to increase as isolated small components as more interactions are included. On the other hand, the CAUS and HYBR predicted interactions are constrained to have one of the known TFs as the source of interactions. Thus, CAUS and HYBR are more likely to grow as a single-component network as more interactions are included.

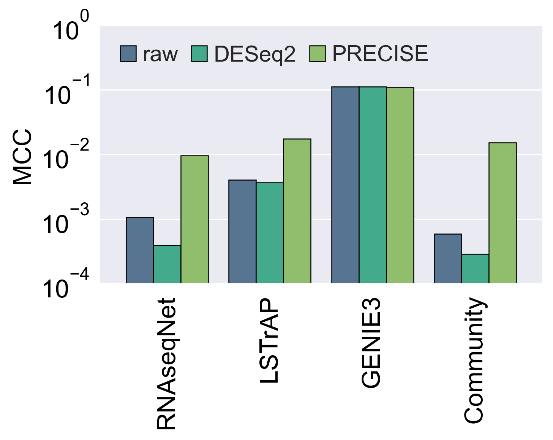

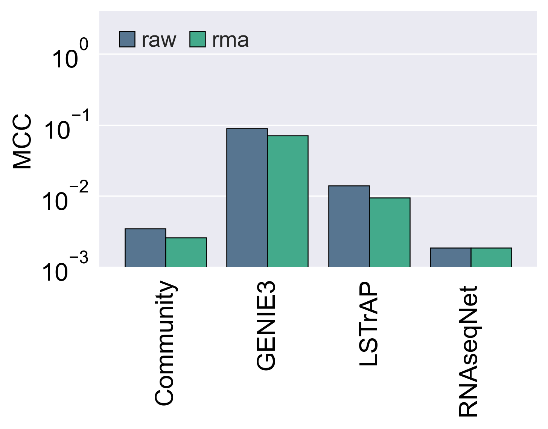
 **A	B**

#### Supplementary Figure 19. Assessment of tools designed to work with RNA-seq data. We used A) RNA-seq and B) Microarray data as input with different normalization levels. The community network was constructed using an only tool designed to work with RNA-seq data (LSTrAP and RNAseqNet). GENIE3 was used for comparison purposes.

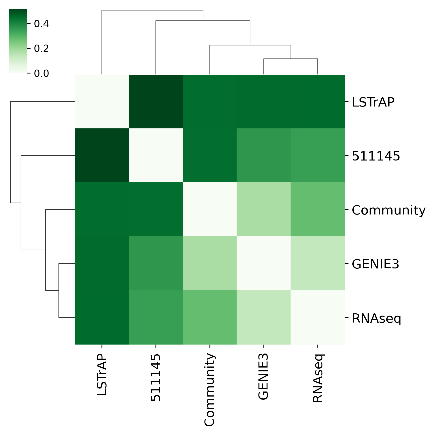

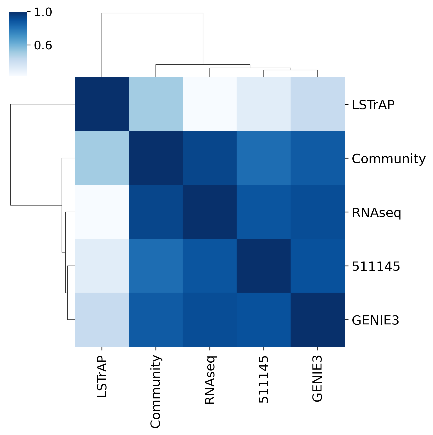

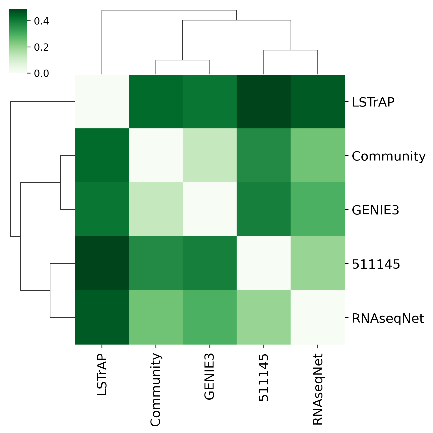

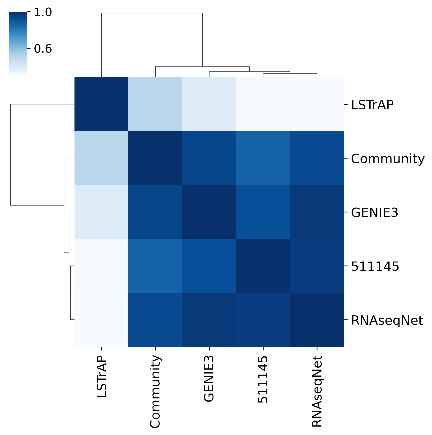

#### Supplementary Figure 20. RNA topology using structural properties and the *D*-value. A) Tools using PRECISE data as input (RNA-seq) and B) using Microarray raw data. We used the raw data for microarray because of the score decline after RMA normalization (Supplementary Figure 18).

### ∑ References

Altay, G., and Emmert-Streib, F. (2010). Inferring the conservative causal core of gene regulatory networks. *BMC Syst Biol* 4**,** 132. doi: 10.1186/1752-0509-4-132.

Bansal, M., Belcastro, V., Ambesi-Impiombato, A., and di Bernardo, D. (2007). How to infer gene networks from expression profiles. *Mol Syst Biol* 3**,** 78. doi: 10.1038/msb4100120.

Bonneau, R., Reiss, D.J., Shannon, P., Facciotti, M., Hood, L., Baliga, N.S., et al. (2006). The Inferelator: an algorithm for learning parsimonious regulatory networks from systems-biology data sets de novo. *Genome Biol* 7(5)**,** R36. doi: 10.1186/gb-2006-7-5-r36.

Budden, D.M., and Crampin, E.J. (2016). Information theoretic approaches for inference of biological networks from continuous-valued data. *BMC Syst Biol* 10(1)**,** 89. doi: 10.1186/s12918-016-0331-y.

Chai, L.E., Loh, S.K., Low, S.T., Mohamad, M.S., Deris, S., and Zakaria, Z. (2014). A review on the computational approaches for gene regulatory network construction. *Comput Biol Med* 48**,** 55-65. doi: 10.1016/j.compbiomed.2014.02.011.

Faith, J.J., Hayete, B., Thaden, J.T., Mogno, I., Wierzbowski, J., Cottarel, G., et al. (2007). Large-scale mapping and validation of Escherichia coli transcriptional regulation from a compendium of expression profiles. *PLoS Biol* 5(1)**,** e8. doi: 10.1371/journal.pbio.0050008.

Haury, A.C., Mordelet, F., Vera-Licona, P., and Vert, J.P. (2012). TIGRESS: Trustful Inference of Gene REgulation using Stability Selection. *BMC Syst Biol* 6**,** 145. doi: 10.1186/1752-0509-6-145.

Hernandez, H. (2018). "Statistical Modeling and Analysis of Experiments without ANOVA".).

Hoffman, J.I.E. (2015). "Chapter 26 - Analysis of Variance II. More Complex Forms," in *Biostatistics for Medical and Biomedical Practitioners,* ed. J.I.E. Hoffman. Academic Press), 421-447.

Huynh-Thu, V.A., Irrthum, A., Wehenkel, L., and Geurts, P. (2010). Inferring regulatory networks from expression data using tree-based methods. *PLoS One* 5(9). doi: 10.1371/journal.pone.0012776.

Kang, Y., Liow, H.H., Maier, E.J., and Brent, M.R. (2018). NetProphet 2.0: mapping transcription factor networks by exploiting scalable data resources. *Bioinformatics* 34(2)**,** 249-257. doi: 10.1093/bioinformatics/btx563.

Kuffner, R., Petri, T., Tavakkolkhah, P., Windhager, L., and Zimmer, R. (2012). Inferring gene regulatory networks by ANOVA. *Bioinformatics* 28(10)**,** 1376-1382. doi: 10.1093/bioinformatics/bts143.

Linde, J., Schulze, S., Henkel, S.G., and Guthke, R. (2015). Data- and knowledge-based modeling of gene regulatory networks: an update. *EXCLI J* 14**,** 346-378. doi: 10.17179/excli2015-168.

Lo, K., Raftery, A.E., Dombek, K.M., Zhu, J., Schadt, E.E., Bumgarner, R.E., et al. (2012). Integrating external biological knowledge in the construction of regulatory networks from time-series expression data. *BMC Syst Biol* 6**,** 101. doi: 10.1186/1752-0509-6-101.

Marbach, D., Costello, J.C., Kuffner, R., Vega, N.M., Prill, R.J., Camacho, D.M., et al. (2012). Wisdom of crowds for robust gene network inference. *Nat Methods* 9(8)**,** 796-804. doi: 10.1038/nmeth.2016.

Margolin, A.A., Nemenman, I., Basso, K., Wiggins, C., Stolovitzky, G., Dalla Favera, R., et al. (2006). ARACNE: an algorithm for the reconstruction of gene regulatory networks in a mammalian cellular context. *BMC Bioinformatics* 7 Suppl 1**,** S7. doi: 10.1186/1471-2105-7-S1-S7.

Proost, S., Krawczyk, A., and Mutwil, M. (2017). LSTrAP: efficiently combining RNA sequencing data into co-expression networks. *BMC Bioinformatics* 18(1)**,** 444. doi: 10.1186/s12859-017-1861-z.

Ruyssinck, J., Huynh-Thu, V.A., Geurts, P., Dhaene, T., Demeester, P., and Saeys, Y. (2014). NIMEFI: gene regulatory network inference using multiple ensemble feature importance algorithms. *PLoS One* 9(3)**,** e92709. doi: 10.1371/journal.pone.0092709.

Young, W.C., Raftery, A.E., and Yeung, K.Y. (2014). Fast Bayesian inference for gene regulatory networks using ScanBMA. *BMC Syst Biol* 8**,** 47. doi: 10.1186/1752-0509-8-47.

Zhang, B., and Horvath, S. (2005). A general framework for weighted gene co-expression network analysis. *Stat Appl Genet Mol Biol* 4**,** Article17. doi: 10.2202/1544-6115.1128.
